# The DEAD-box RNA helicases RhlE2 is a global regulator of *Pseudomonas aeruginosa* lifestyle and pathogenesis

**DOI:** 10.1101/2021.01.29.428592

**Authors:** Stéphane Hausmann, Diego Gonzalez, Johan Geiser, Martina Valentini

## Abstract

The RhlE DEAD-box RNA helicase protein family is widespread among Proteobacteria, but it is the least understood due to the lack of a clear biological function. Here, we study the two RhlE homologs present in the opportunistic pathogen *Pseudomonas aeruginosa*. RhlE1 and RhlE2 diverged during *P. aeruginosa* evolution; our data indicate that this resulted in a non-redundant biological role, a distinct molecular action and an enzymatic activity differentially stimulated by RNA. Whereas RhlE1 is specifically necessary for bacteria growth in cold, we show that RhlE2 acts as global post-transcriptional regulator, affecting the level of hundreds of cellular transcripts and multiple functionalities indispensable not only for *P. aeruginosa* environmental adaptation, but also for its virulence. The global action of RhlE2 relies on a unique C-terminal extension, which establishes an RNA-dependent interaction with the RNase E endonuclease and the cellular RNA degradation machinery.

## Introduction

DEAD-box RNA helicases are RNA-binding proteins that upon ATP binding and hydrolysis remodel RNA structure and RNA–protein complexes (1). DEAD-box RNA helicases are found in all three domains of life, as well as in some viruses, and they are identified on the basis of highly conserved amino acid motifs within the catalytic core, “D-E-A-D” being one of those motifs (2,3). Besides the conserved catalytic region, they possess variable N- and C-terminal extensions that can regulate the core catalytic activity or coordinate interactions with RNA molecules or proteins (4–7). The *Escherichia coli* DbpA RNA helicase, for example, contains an RNA-binding domain (RBD) at the C-terminus, which is responsible for a specific and tight binding to hairpin 92 of 23S ribosomal RNA (8).

From bacteria to humans, DEAD-box RNA helicases are involved in maintaining a correct basal RNA metabolism, by controlling mRNA translation, RNA degradation and ribosome biogenesis (1). In bacteria, though, DEAD-box RNA helicases do not perform only housekeeping functions, but can also regulate their environmental adaptation, host colonization and, in case of pathogens, infectious processes (9–11).

Bacterial DEAD-box RNA helicases can be classified into to three major phylogenetic groups, that are distinguished by the presence of DbpA-RBD: DbpA RBD-containing proteins, DbpA RBD-lacking proteins and RhlE-like proteins (12). The RhlE-like RNA helicases, which are named from the *E. coli* DEAD-box RNA helicase belonging to this group, seem to have originated from a duplication of a DbpA-RBD lacking protein in a Proteobacterial ancestor and have successively expanded in this phylum, where they can be present in several copies per genome (12). The role and molecular action of RhlE-like proteins are not understood, and they vary across species. No phenotypic defect linked to *rhlE* deletion has been described in *E. coli*, *Pseudomonas syringae* LZ4W and *Mycobacterium tuberculosis*, while in *Yersinia pseudotuberculosis* and *Caulobacter crescentus* deletion of *rhlE* affects cold acclimation (13–18). In *E. coli*, RhlE seems to liaise with the SrmB and CsdA RNA helicases during ribosome biogenesis, although the exact molecular action is not yet deciphered, and to interact with the RNase E endonuclease for RNA degradation, even if the conditions in which the interaction occurs in vivo are unclear (19,20). In *P. syringae* LZ4W, *M. tuberculosis* and *C. crescentus*, RhlE is described as a core component of the cellular RNA degradosome machinery, hence involved in RNA degradation (13,17,18).

In the present study, we investigate the role of RhlE proteins in the Gram-negative bacterium *Pseudomonas aeruginosa*. The bacterium has a remarkable distribution across multiple ecological niches, including a wide variety of hosts (21–23). In humans, *P. aeruginosa* infections pose a significant life threat to immunocompromised and immunodeficient persons, intensive care patients, patients with cystic fibrosis or other forms of bronchiectasis and chronic obstructive pulmonary diseases (24–28). As opportunistic pathogen, *P. aeruginosa* possesses a remarkable virulence plasticity, being able to colonize different type of organs and modify its virulence strategy and physiology in response to the host context (29). Scarce information exists on *P. aeruginosa* DEAD-box RNA helicase genes and their role on the regulating the bacterium versatility (30). In addition, we noticed that the genome of *P. aeruginosa* encodes two *rhlE* genes: PA3950 (*rhlE1*) and PA0428 (*rhlE2*), making the bacterium an ideal model to assess any redundancy and neofunctionalization among duplicated RhlE proteins. Here, we reveal that both RhlE1 and RhlE2 are important for *P. aeruginosa* cold adaptation, but *rhlE2* mutation results in a broader pleiotropic effect on motility, biofilm formation and virulence. We assess RhlE1 and RhlE2 RNA preferences in vitro and we identify the RhlE2 regulon in vivo. Moreover, we define a model of RhlE2 mechanism of action whereby RhlE2 affects RNA stability via an unusual interaction with RNase E, which is dependent on RNA and RhlE2 C-terminal extension. Finally, we discuss the phylogenetic diversity of RhlE proteins by comparing *P. aeruginosa* RhlE proteins with previously characterized RhlE proteins in other Proteobacteria.

## Results

### *Pseudomonas aeruginosa* encodes two RhlE-like RNA helicases with different origin and features

The RhlE-like group RNA helicases is widespread in Proteobacteria, with sometimes several homologs present in a single genome (9,12). In most *Pseudomonas* species two RhlE homologs are found, belonging to two different terminal branches of the RhlE phylogenetic tree that groups eight main clades (**Fig 1A** and **S1**). In *P. aeruginosa* PAO1 genome, two genes encode for RhlE-like proteins, PA3950 (RhlE1) and PA0428 (RhlE2), having 51% and 43% sequence identity to the *E. coli* RhlE, respectively (**Fig 1B**). RhlE1 is found on a branch represented mostly in environmental bacteria, including *Vibrio, Shewanella, Legionella* or *Azotobacter* species; while the clade where the RhlE2 belongs includes many *Pseudomonas* genus (**Fig S1**). Alignment of representative RhlE homologs shows that the canonical RecA1 and RecA2 RNA helicase domains are conserved, including the ten short motifs typical of DEAD-box RNA helicases, while high sequence divergence is present at the C-terminus of the proteins (**Fig S2**). Clustering of the C-terminal extension among RhlE homologs does not show any global pattern; it is therefore likely that C-terminal regions are clade-specific and fast evolving. In *P. aeruginosa*, RhlE1 possesses a short CTE that contains a stretch of lysine-rich amino acids, while RhlE2 possess the longest CTE among RhlE-like proteins that includes a stretch of ~50 amino acids rich in glycine-glutamate (**Fig S3**).

**Figure 1.**
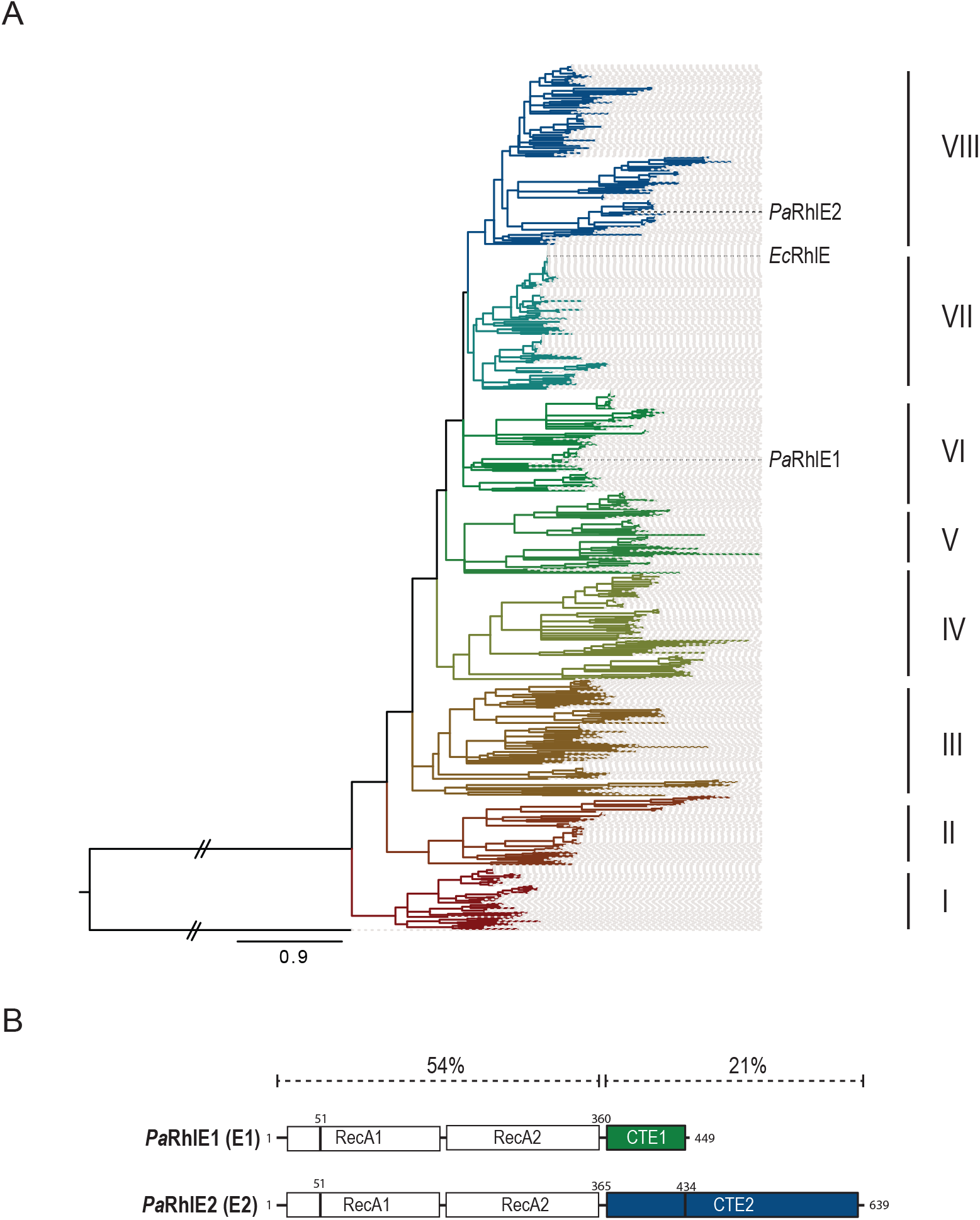
Phylogenetic tree of the bacterial RhlE DEAD-box proteins. **(A)** The tree is based on 799 sequences of bacterial DEAD-box proteins, found in the NCBI RefSeq database. The *P. aeruginosa* RecQ protein (PA3344) has been used to root the tree. At least 8 distinct lineages can readily be identified and are divided into group I to VIII. **(B)** Linear representation of *P. aeruginosa* (Pa) RhlE1 (E1, PA3950) and RhlE2 (E2, PA0428) with the RecA1 and RecA2 domains in white boxes and their C-terminal extension (CTE) in green and blue boxes, respectively. % indicates sequence identity with *E. coli* RhlE protein. Lines corresponds to recombinantly produced RhlE1 and RhlE2 variants (K51A and 1-434)

### RhlE1 and RhlE2 are both necessary for *P. aeruginosa* cold adaptation, but are not redundant

Cold sensitivity is a phenotype that has been previously associated to inactivation of RNA helicase genes and in particular to *rhlE* inactivation in *Y. pseudotuberculosis* and *C. crescentus* (9,13,14). To investigate the role of RhlE homologs in *P. aeruginosa*, we constructed single *rhlE1* and *rhlE2* deleted strains as well as a double Δ*rhlE1*Δ*rhlE2* mutant and we tested growth on cold. Specifically, we evaluated the growth properties of the deletion strains compared to the wild type by spotting serial dilutions of cells (from liquid cultures adjusted to an OD600 of 1.0) on LB agar medium and incubating plates at 37°C and 16°C. At 37°C all mutants displayed similar growth as the wild type strain, as shown by CFU counting of serial dilutions (**Fig 2A**), as well as by OD measurements of liquid cultures at different time points (**Fig S4**). By contrast, at 16°C, deletion of *rhlE1* or *rhlE2* resulted in a growth defect, which was further increased in the Δ*rhlE1*Δ*rhlE2* mutant (**Fig 2B**). Both RhlE1 and RhlE2 are shown to be important for cold adaptation, but while loss of *rhlE1* produced a severe growth delay, loss of *rhlE2* was only slightly deleterious. We then tested if the functions of RhlE1 and RhlE2 overlap during cold growth, by preforming complementation experiments. To do so, we employed the mini-Tn7 system to insert into the chromosomal Tn7 attachment site of Δ*rhlE1*, Δ*rhlE2* or Δ*rhlE1*Δ*rhlE2* mutants a copy of *rhlE1* or *rhlE2* gene under the control of an arabinose-inducible promoter. As the genes carried by the mini-Tn7 constructs were cloned in frame with a N-terminal 3xFLAG epitope, we could verify their expression levels by Western blot, which were comparable (**Fig S5**). As shown in **Fig 2C**, growth of the *rhlE1* mutant at 16°C could be restored to wild type levels by the insertion of the mini-Tn7_3xFLAG-RhlE1 (∷3xFLAG-E1) construct, likewise for the complementation of *rhlE2* mutant by the mini-Tn7_3xFLAG-RhlE2 (∷3xFLAG-E2). However, it was not possible to fully restore the growth at 16°C of the Δ*rhlE1*Δ*rhlE2* mutant by reintroduction *in trans* of only one *rhlE* gene **(Fig 2C)**, nor to complement the phenotype of the *rhlE1* mutant by 3xFLAG-E2 and vice versa (**Fig S6**). Altogether, these data provide evidence that RhlE1 and RhlE2 perform unique and specific regulatory actions during growth on cold, as they are not redundant and do not cross-complement each other.

**Figure 2.**
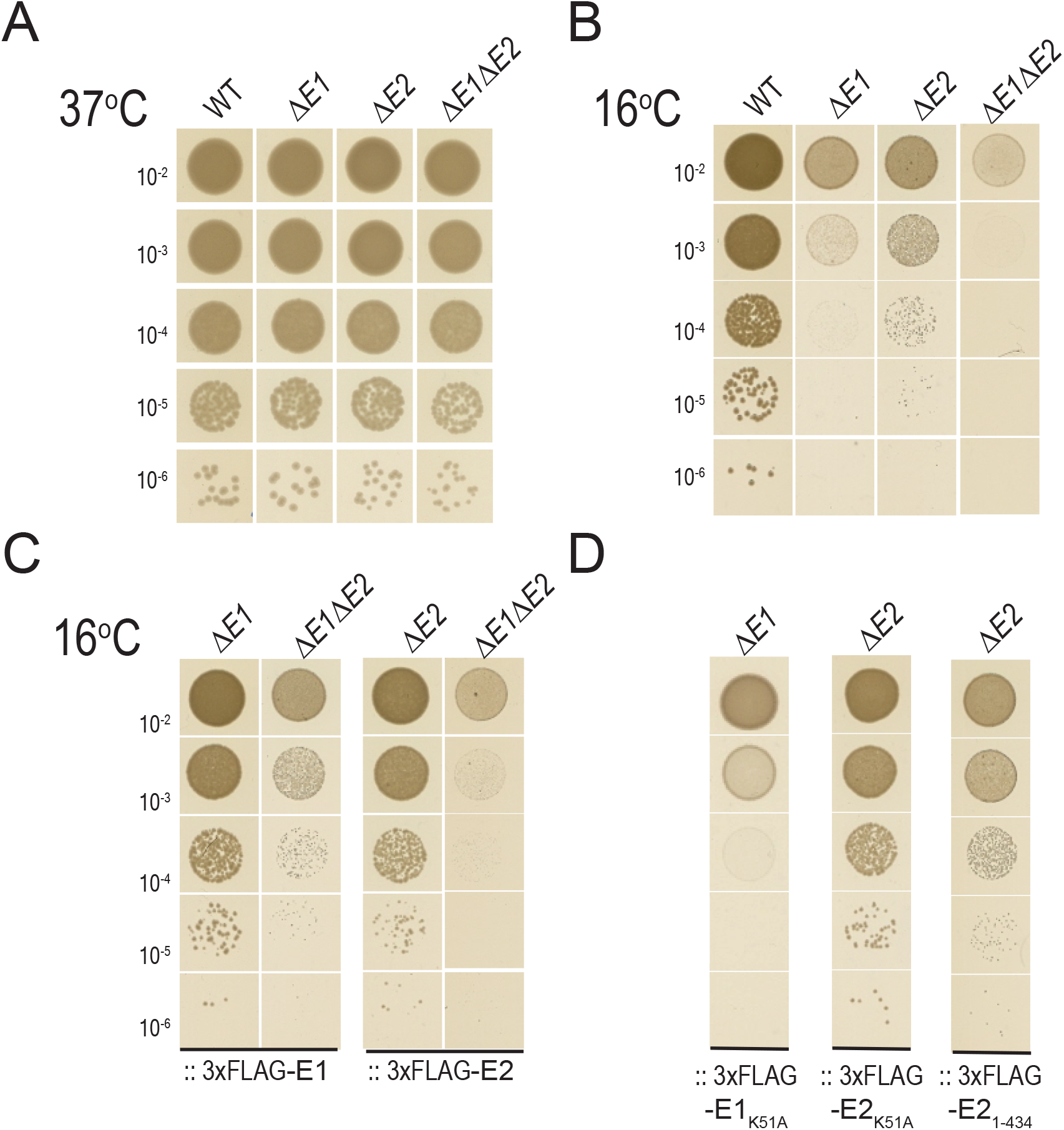
Role of RhlE1 and RhlE2 in *P. aeruginosa* cold adaptation. **(A)** 37 °C and **(B)** cold shock (16 °C) survival assays of wild type PAO1, Δ*rhlE1* mutant (ΔE1), Δ*rhlE2* mutant (ΔE2) and Δ*rhlE1*Δ*rhlE2* mutant (ΔE1ΔE2). **(C)** Complementation effects of cold (16 °C) growth using mini-Tn7 constructs expressing 3xFLAG-tagged RhlE1 (∷3xFLAG-E1), 3xFLAG-tagged RhlE2 (∷3xFLAG-E2), **(D)**, 3xFLAG-tagged RhlE1_K51A_ (∷3xFLAG-E1_K51A_), 3xFLAG-tagged RhlE2_1-434_ (∷3xFLAG-E2_1-434_) and 3xFLAG-tagged RhlE2_K51A_ (∷3xFLAG-E2_K51A_). Strains background is indicated, and experimental details are described in Materials and Methods. At least three independent biological replicates were tested, results from a representative plate is shown.

### RhlE2 is a global lifestyle regulator in *P. aeruginosa*

Some bacterial RNA helicases have been shown to regulate other cell functions besides cold growth and to be involved in environmental adaptation or even pathogenicity (9,11). Thus, we tested whether RhlE1 and RhlE2 could also impact medically relevant physiology and behaviour of *P. aeruginosa*. We first tested the *rhlE* mutant strains motility in liquid and on surfaces (**Fig 3A**), then their capacity to form early and mature biofilms (**Fig 3B**). Having ruled out a growth defect of the strains at 37°C, all the experiments were carried at this temperature. Interestingly, the Δ*rhlE2* mutant was strongly affected on all these traits, being less motile (20, 2.5- and 15-fold reduction of swimming, twitching and swarming, respectively) and significantly impaired in biofilm formation (3- and 1.5-fold decrease of early and mature biofilm, respectively) when compared to the wild type strain. By contrast, deletion of *rhlE1* did not results in significant differences compared to wild type, while the Δ*rhlE1*Δ*rhlE2* mutant phenotype didn’t differ from the Δ*rhlE2* strain. Altogether, these results revealed that RhlE2 affects several functionalities and cellular processes in *P. aeruginosa*, thus acting as global lifestyle regulator; while RhlE1 role is specific to cold adaptation. This conclusion is validated by the observation that complementation of the Δ*rhlE1*Δ*rhlE2* mutant with 3xFLAG-E2 was able to fully restore swarming motility to wild type levels, whereas the Tn7 construct carrying *rhlE1* failed (**Fig 3C**). Finally, the lack of a phenotype linked to a *rhlE1* deletion even in absence of *rhlE2* indicates that no buffering exists among these two RNA helicases **(Fig. 3A and 3B).**

**Figure 3.**
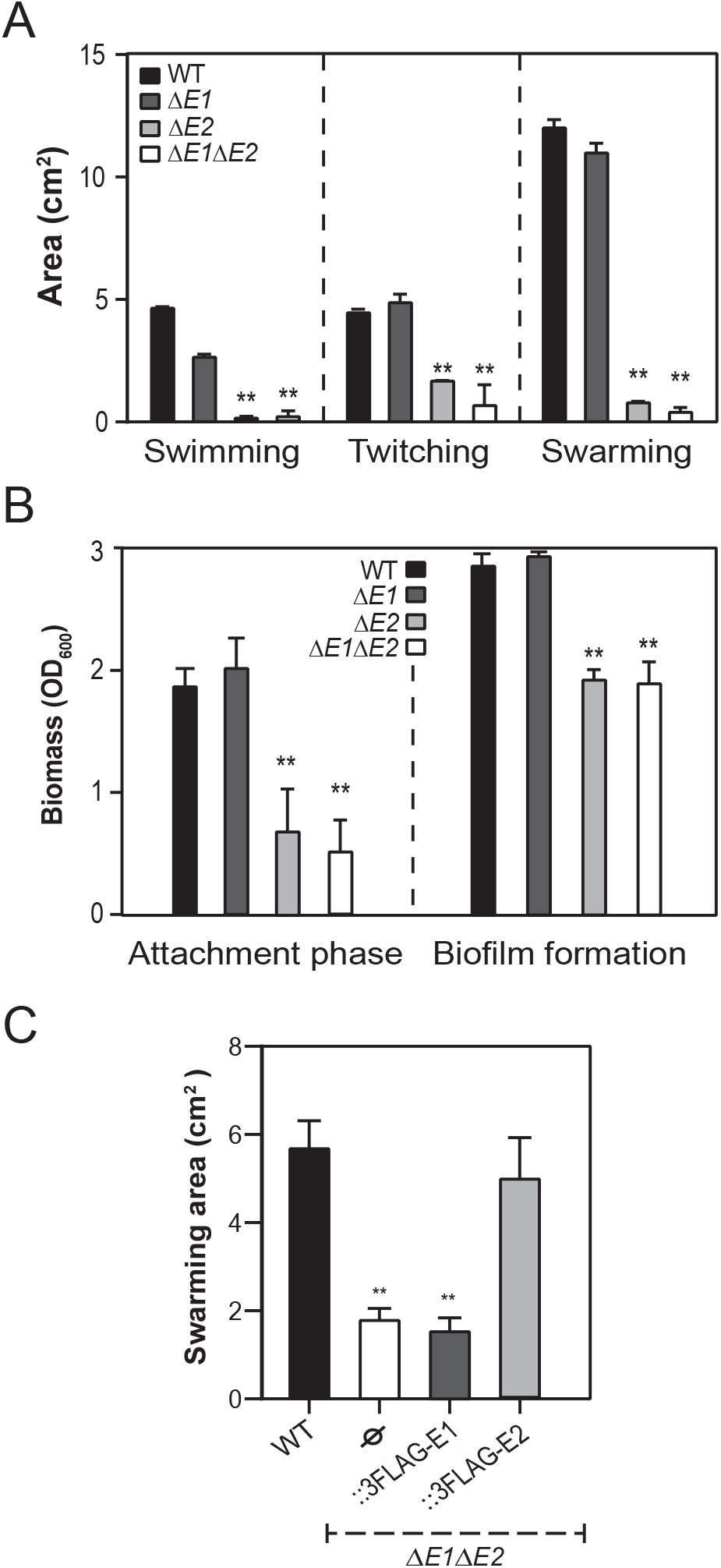
Role of RhlE1 and RhlE2 in *P. aeruginosa* motilities and biofilm formation. **(A)** Surface area covered by the swimming, swarming and twitching wild type PAO1 (WT, black diamonds), the Δ*rhlE1* mutant (ΔE1,dark grey triangles), the Δ*rhlE2* mutant (ΔE2, light grey circles) and the Δ*rhlE1*Δ*rhlE2* mutant (ΔE1ΔE2, white squares) cells (± standard deviation) as calculated by averaging data from four individual plates. **(B)** Surface attached biomass, measured by crystal violet staining, of wild type PAO1 and *rhlE1/rhlE2* mutant strains after 6 hours (attachment phase) or 12 hours (biofilm formation) of growth at 37°C in 24-microtiter plates. **(C)** Complementation of swarming motility using mini-Tn7 constructs carrying 3xFLAG-tagged RhlE1 (∷3xFLAG-E1), 3xFLAG-tagged RhlE2 (∷3xFLAG-E2) expressed in a Δ*rhlE1*Δ*rhlE2* mutant Each value is the average of three different cultures ± standard deviation. (*, *p* < 0.05; **, *p* < 0.01)

### RNA sequencing analysis reveals the importance of RhlE2 for *P. aeruginosa* virulence

To examine further the role of RhlE2 in *P. aeruginosa*, we compared the transcriptome of *rhlE2* deleted strains in the wild type or in the Δ*rhlE1* background, by performing RNA-sequencing. As experimental condition, we choose to investigate the role of RhlE2 in swarming cells (**Fig 4A**). Swarming motility is a coordinated movement of a bacterial population on semi-solid surfaces (0.5 to 0.7% agar), which is mediated by flagella rotation and surfactants secretion (31). Importantly, *P. aeruginosa* swarming cells possess a physiological state which is highly relevant to the bacterium pathogenicity as they express virulence factors, they perform epithelial surfaces colonization and are associated to the establishment of lungs infection through biofilm formation (32,33). Approximately 15% of the transcriptome was significantly affected by *rhlE2* mutation when compared to wild type PAO1 (N=836 genes, *p*-value <0.05, FC>2), of which 4% (N=228) of transcripts were up-regulated and 11% (N=608) down-regulated (**Fig 4B**, **Table S1**). Differentially regulated genes were grouped into the corresponding KEGG pathway: most significantly downregulated genes code for ribosomal proteins, metabolic pathways and enzymes involved in the biosynthesis of secondary metabolites, while upregulated ones are linked to xenobiotics biodegradation and metabolism (e.g. 4-fluorobenzoate, chlorocyclohexane and chlorobenzene degradation, **Table S2**). Consistent with the phenotypic analysis, *rhlE1* deletion had not significant impact on the cellular transcript levels neither in the wild type nor in the *rhlE2* mutant background (**Fig 4B**). Next, we validated the transcriptome analysis by choosing a set of putative RhlE2 targets and examining their abundance by RT-qPCR in the Δ*rhlE1*, Δ*rhlE2* and Δ*rhlE1*Δ*rhlE2* mutants as compared to the wild type, starting from cell grown under experimental conditions similar to those used for the original RNA-sequencing analysis. The gene chosen have been previously shown to either regulate swarming or being regulated in this condition, such as *roeA*, coding for a diguanylate cyclase (34), *lhpF,* coding for the metabolic enzyme D-hydroxyproline dehydrogenase (35), the *lasB* elastase (36) and the *pvdF* pyoverdine synthase (37) encoding genes, and finally the *rhlA* gene was chosen as control (rhamnosyltransferase chain A) being unaffected by *rhlE2* mutation, but previously shown necessary for swarming motility (38). In agreement with the RNA-seq data, *lasB* and *pvdF* mRNA levels were, respectively, 10.25 and 10.41-fold down-regulated in the Δ*rhlE2* mutant when compared to the wild type; while *roeA* and *lhpF* were, respectively, 3.98 and 8.94-fold up-regulated and *rhlA* mRNA levels were unchanged (**Fig 4C**). To further validate the observation concerning regulation of *lasB* expression, a transcriptional/translational *lasB’-‘lacZ* fusion (39) was assayed for β-galactosidase activity in the four strains, at different growth phases in NYB (**Fig 4D**). Expression of *lasB* is dependent on cell-density and therefore maximal at stationary phase (40), the same trend was observed in all mutants but in the Δ*rhlE2* and Δ*rhlE1*Δ*rhlE2* mutants the expression of the *lasB’-‘lacZ* fusion was reduced of ~2.5-fold in stationary phase, when compared to the wild type and Δ*rhlE1* strains.

**Figure 4.**
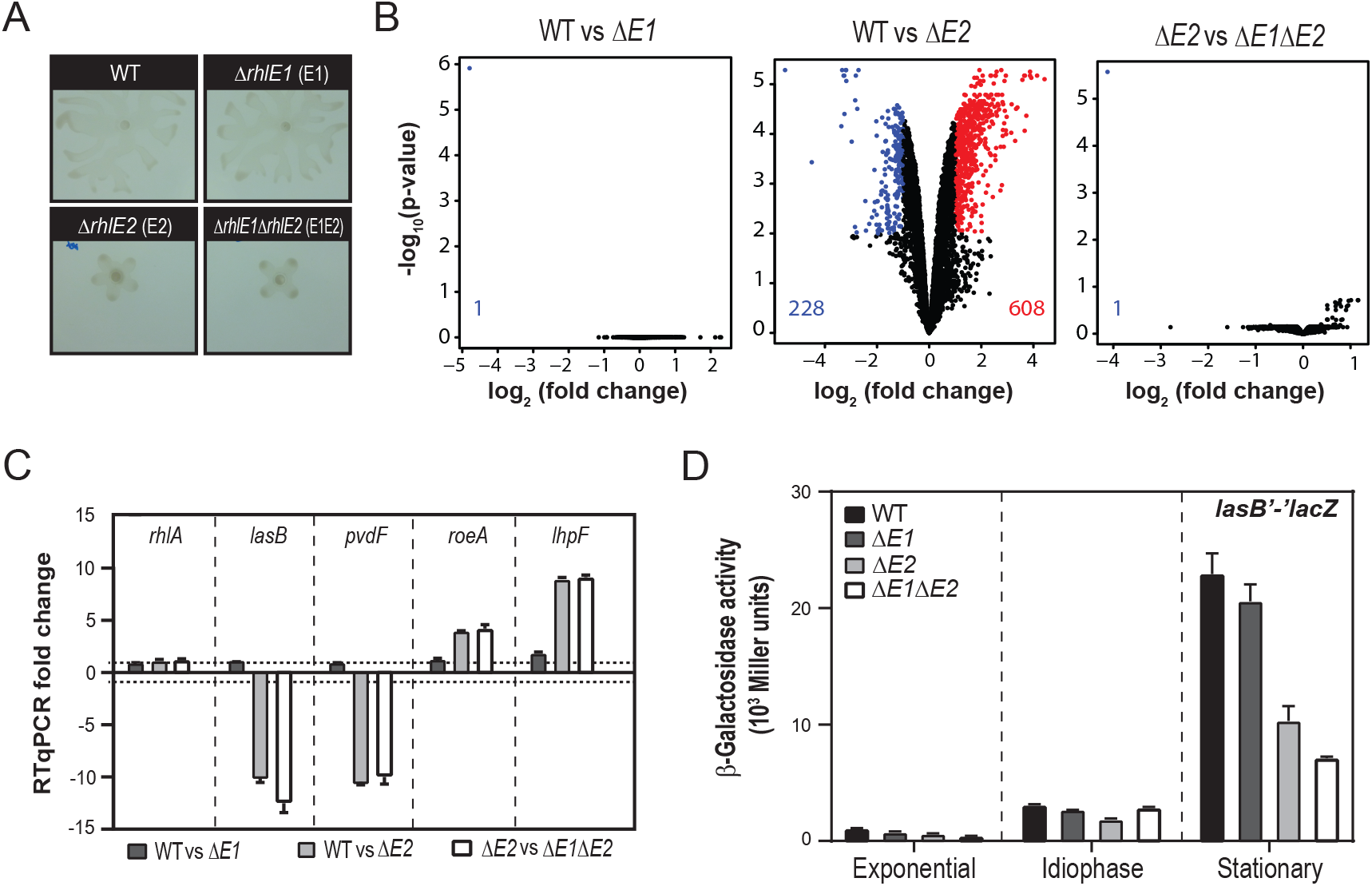
The RhlE2 regulon. **(A)** Example of wild type PAO1, Δ*rhlE1* mutant, Δ*rhlE2* mutant and the Δ*rhlE1*Δ*rhlE2* mutant swarming cells that were subjected to RNA-sequencing analysis. **(B)** Volcano plot representation of transcriptome comparison between the wild type PAO1 (WT) and Δ*rhlE1* (ΔE1) or Δ*rhlE2* (ΔE2) or Δ*rhlE1*Δ*rhlE2* (ΔE1ΔE2). The genes are colored black if they pass the thresholds for −log10 P value (P value < 0.05) and log fold change |FC| ≥2. **(C)** Relative transcript abundance in the wild type PAO1 and *rhlE1/rhlE2* mutant strains as determined by RT-qPCR (RT-qPCR fold change). Values are represented as averages of 3 independent replicates for every strain. **(D)** Cell density-dependent β-galactosidase expression of a transcriptional/translational *lasB’-‘lacZ* fusion carried by plasmid pTS400 (39) in wild type PAO1 (black bars), Δ*rhlE1* mutant (ΔE1,dark grey bars), Δ*rhlE2* mutant (ΔE2, light grey bars) and Δ*rhlE1*Δ*rhlE2* mutant (ΔE1ΔE2, white bars). Each value is the average of three different cultures ± standard deviation.

Besides *lasB*, one interesting observation concerning the *rhlE2* mutant transcriptome was the significant downregulation of various other genes encoding virulence factors (**Table S1**). To assess whether mRNA levels regulation by RhlE2 also results in changes of virulence factor abundance, we measured in our strains the extracellular levels of the LasB elastase and pyocyanin, a redox-active molecule inducing oxidative stress in host cells (41). We found that the Δ*rhlE2* and Δ*rhlE1*Δ*rhlE2* mutants produced 2.6- and 3.6-fold less elastase and 2,1- and 2.4-fold less pyocyanin than the wild type, respectively, while the *rhlE1* deletion did not produce any significant difference (**Fig 5A-B**).

**Figure 5.**
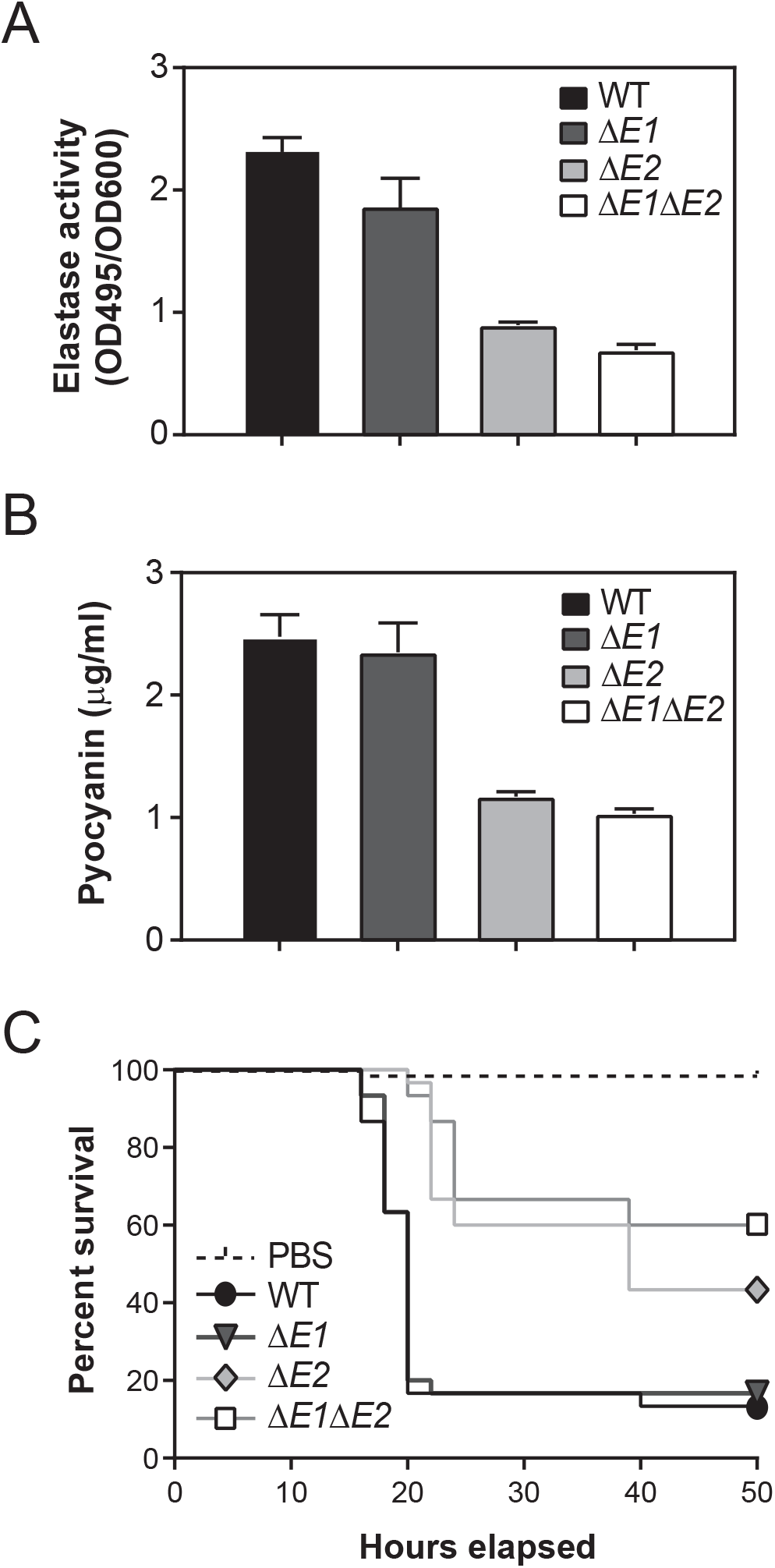
RhlE2 regulation of *P. aeruginosa* virulence. **(A)** Elastase activity and **(B)** pyocyanin levels of wild type PAO1 (black bars), Δ*rhlE1* mutant (ΔE1, dark grey bars), Δ*rhlE2* mutant (ΔE2, light grey bars) and Δ*rhlE1*Δ*rhlE2* mutant (ΔE1ΔE2, white bars) supernatant. **(C)** Survival curves of *G. mellonella* larvae infected with wild type PAO1 and *rhlE1/rhlE2* mutant strains. PBS, larvae injected with sterile physiological solution. Each experiment (A-C) was repeated at least with three different independent cultures for each strain, values indicated average from biological replicates ± standard deviation. For experimental details see Material and Methods.

### RhlE2 affects *P. aeruginosa* virulence in *Galleria mellonella*

*Galleria mellonella* has become an important model to study bacterial pathogenesis (42). We used this system to confirm that RhlE2 has indeed an important role during *P. aeruginosa* pathogenesis *in vivo*. In agreement with data disclosed above, the Δ*rhlE2* and Δ*rhlE1*Δ*rhlE2* mutants displayed a significant attenuation of virulence in the *G. mellonella* infection model. Indeed, at 24 hours post-infection the mortality of larvae injected with either Δ*rhlE2* or Δ*rhlE1*Δ*rhlE2* reached less than 40%, while the mortality rate of larvae challenged with wild type or Δ*rhlE1* strain was 80% (**Fig 5C**). Altogether, these results indicate that RhlE2 controls, directly and/or indirectly, the expression of virulence or virulence-related genes and is necessary for *P. aeruginosa* pathogenicity *in vivo*.

### RhlE1 and RhlE2 display an RNA-dependent ATPase activity in vitro

To fuel their molecular actions, RNA helicases possess an ATP hydrolytic activity that is dependent on RNA binding (2). To confirm this prediction, based on RhlE1 and RhlE2 sequence conservation, we assess their enzymatic activity in vitro. The proteins were produced in *E. coli* as N-terminal His^10^-Smt3-tagged fusion and purified from a soluble extract by adsorption to nickel-agarose and elution with 250 mM imidazole (**Fig 6A**). As control, we also purified the RhlE1 and RhlE2 catalytic mutants, in which the Lys51 of RhlE1 and RhlE2 within motif I (involved in ATP binding and hydrolysis) is replaced by Ala, making the proteins inactive (43). As shown in **Fig 6B**, recombinant RhlE1 and RhlE2 catalyzed the release of ^32^Pi from [*γ*-^32^P] ATP in the presence of poly(U) RNA and that the extent of ATP hydrolysis was proportional to the protein concentration. From the slope of the titration curves, we estimated that RhlE1 and RhlE2 hydrolyzed 0.0119 nmol and 0.0068 nmol ATP per ng of enzyme, respectively, during the 15 min reaction. These values translate into a turnover number of 51 min^−1^ for RhlE1 and 37 min^−1^ for RhlE2 indicating that both RhlE proteins have a similar ATPase activity in the presence of poly(U) RNA. As expected for DEAD-box RNA helicases, both RhlE1 and RhlE2 catalyzed no detectable ATP hydrolysis in absence of RNA (**Fig 6B**). Moreover, the ATPase activity of RhlE1_K51A_ and RhlE2_K51A_ mutants were less than 1% of the activity of the respective wild type enzyme stimulated by poly(U) RNA (**Fig 6C**), validating that the observed RNA dependent phosphohydrolase activities of wild type RhlE1 and RhlE2 are intrinsic to the recombinant proteins.

**Figure 6.**
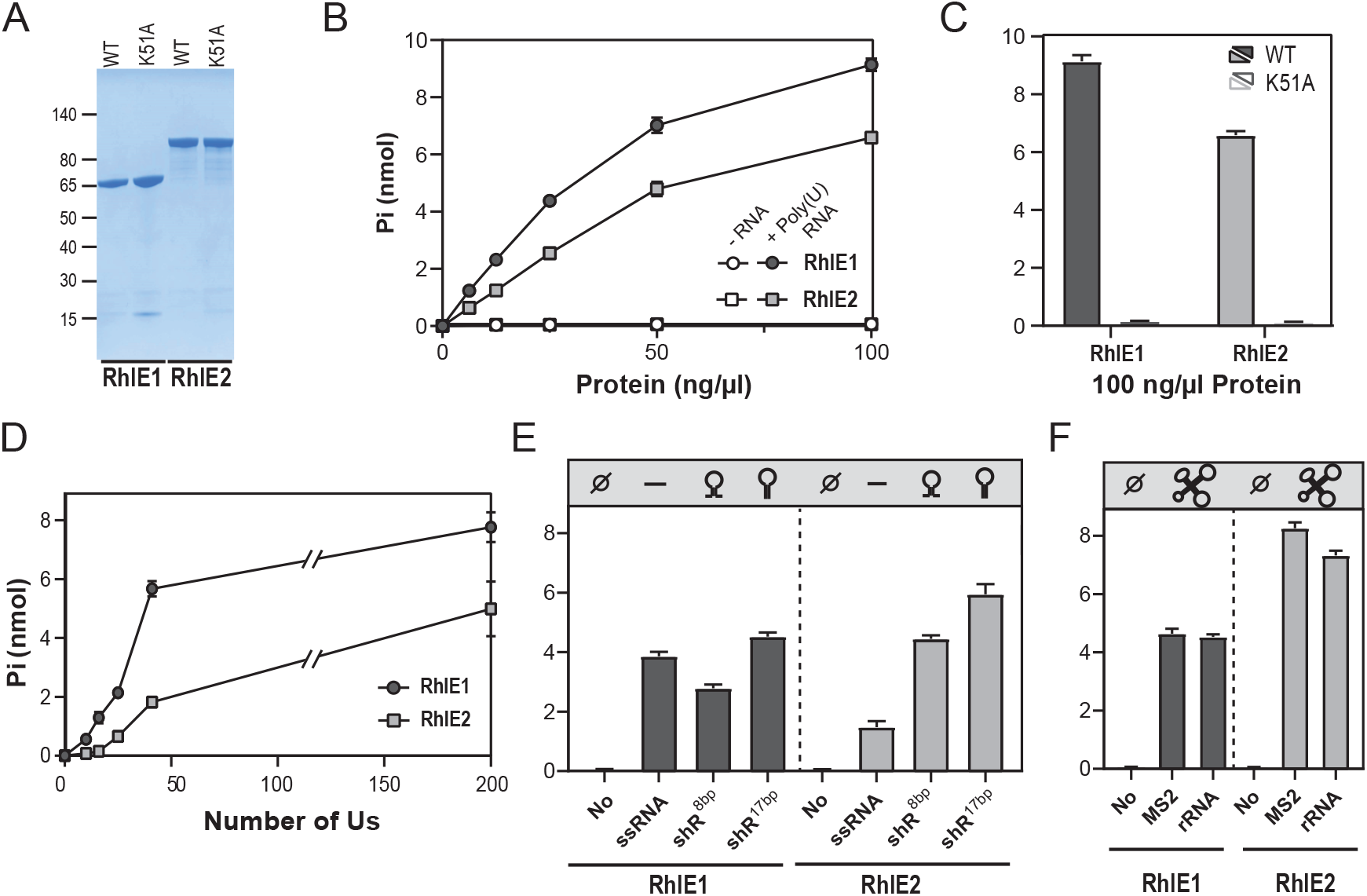
ATPase activity of recombinant RhlE1 and RhlE2. **(A)** RhlE proteins purification. Aliquots (2.5 μg) of the nickel-agarose preparations of wild type (WT) RhlE1(lane 1), and mutant K51A (lane 2), wild type (WT) RhlE2 (lane 3), and mutant K51A (lane 4) were analyzed by SDS-PAGE. The polypeptides were visualized by staining with Coomassie Blue dye. The positions and sizes (kDa) of marker polypeptides are indicated on the left. **(B)** RhlE proteins ATPase activity. Reaction mixtures (15 μl) containing 50 mM Tris-HCl (pH 8.0), 5 mM DTT, 2 mM MgCl2, 1 mM [*γ*-^32^P] ATP, 250 ng/μl Poly(U) (or no RNA; empty symbol), and enzyme as specified were incubated for 15 min at 37 °C. Pi release was determined as described in Materials and Methods and was plotted as a function of input protein. **(C)** RhlE proteins catalytic mutants. Reaction mixtures (15 μl) containing 50 mM Tris-HCl (pH 8.0), 5 mM DTT, 2 mM MgCl2, 1 mM [*γ*-^32^P] ATP, 250 ng/μl Poly(U), and 100 ng/μl of enzyme as specified were incubated for 15 min at 37 °C. The extends of ATP hydrolysis are plotted. **(D)** ATPase activity and RNA length. Reaction mixtures (15 μl) 50 mM Tris-HCl (pH 8.0), 5 mM DTT, 2 mM MgCl2, 1 mM [*γ*-^32^P] ATP, 2 μM polyribouridylic acid (either U41, U26, U16 or U10) as specified, and 50 ng/μl of RhlEs were incubated for 15 min at 37 °C. The extends of ATP hydrolysis are plotted. **(E)** RNA hairpin recognition. Reaction mixtures (15 μl) 50 mM Tris-HCl (pH 8.0), 5 mM DTT, 2 mM MgCl2, 1 mM [*γ*-^32^P] ATP, 2 μM of an oligoribonucleotide as specified and 50 ng/μl of RhlEs were incubated for 15 min at 37 °C. The extends of ATP hydrolysis are plotted. **(F)** Recognition of structured RNA. Reaction mixtures (15 μl) 50 mM Tris-HCl (pH 8.0), 5 mM DTT, 2 mM MgCl2, 1 mM [*γ*-^32^P] ATP, 80 ng/μl MS2 or 100 ng/μl *E. Coli* rRNA of and 50 ng/μl of RhlEs were incubated for 15 min at 37 °C. The extends of ATP hydrolysis are plotted. (B-F) Data are the average ± SEMs from three independent experiments.

### Characterization of the RhlE1 and RhlE2 RNA preferences

As the size of poly(U) RNA is heterogeneous (> 150 ribonucleotides), we examined the effect of RNA length on the ATPase activity of RhlE1 and RhlE2 by using a set of polyribouridylic acid of defined length (U10, U16, U25 and U41, respectively). We found that the extent of phosphate release by RhlE1 with U41, U25, U16, U10 RNA substrate were 73, 27, 16, and 6% of the activity with poly(U) RNA, respectively, while the corresponding RhlE2 ATPase activity was 35, 12, 1 and <1 % (**Fig 6D**). This result indicates that RhlE2 ATPase activity requires RNA oligomers of ≥ 25 nucleotides (nt), whereas RhlE1 can still be activated by RNA oligomers shorter than 16 nt.

We next examined whether the RNA secondary structure can affect the ATPase activity of RhlE proteins. Therefore, we used three different 41-mer RNA substrates: a single-strand RNA (ssRNA), a stem loop of 8 base-pairs with a 9-Us extension both at the 5’ and 3’ end (shRNA^8bp^), and a blunt-ended stem-loop RNAs with 17 base pairs in the stem region (shRNA^17bp^; see **Fig S7**). The phosphate release by RhlE1 was similar between the three different RNA substrates, whereas the ATPase activity of RhlE2 was increased 3-fold with shRNA^8bp^ and 4-fold with a blunt-ended stem-loop RNA shRNA^8bp^, compared to single strand RNA (ssRNA), indicating that RNA structure affects significantly the ATPase activity of RhlE2 but not of RhlE1 (**Fig 6E**). Besides, similar release of phosphate by RhlE1 and RhlE2 is observed with the two 41-mer U41 and ssRNA single-strand RNAs (5.7 nmol and 3.9 nmol Pi for RhlE1 and 1.8 nmol and 1.5 nmol Pi for RhlE2, respectively), confirming that *in vitro* ATPase of DEAD-box helicases display poor RNA sequence specificity (44–46). We also examined the stimulatory effect of naturally occurring long-structured RNAs, namely MS2 RNA and *E. coli* rRNA. We found that MS2 RNA and *E. coli* rRNA are strong activators of RhlE2 ATPase activity than poly(U) RNA, displaying respectively 172% and 153% of the activity measured with poly(U); whereas RhlE1 ATPase activity was 66% and 62% of the activity with poly(U), respectively (**Fig 6F**). Altogether, these results strongly suggest that ATPase activity of RhlE1 and RhlE2 are differently stimulated depending on the length and structure of the RNA.

### The ATPase activity of RhlE1 is necessary for cold adaptation while the RhlE2 ATPase activity is sometimes dispensable

Having validated that the replacement of the lysine by alanine in the motif I abolished the catalytic activity of RhlE1 and RhlE2, we sought to determine if the observed catalytic activity of RhlE1 or RhlE2 was necessary for their role *in vivo*. We constructed mini-Tn7 variants expressing the 3xFLAG-RhlE1_K51A_ or 3xFLAG-RhlE2_K51A_ mutant and we assessed their capacity to complement the phenotypes of the corresponding deleted strain. Expression of the RhlE1 catalytic mutant (3xFLAG-E1_K51A_) was not able to restore the growth at 16°C of the *rhlE1* mutant, while surprisingly the RhlE2_K51A_ construct (3xFLAG-E2_K51A_) was able to sustain growth at 16°C of the Δ*rhlE2* mutant (**Fig 2D**). We checked that growth of the Δ*rhlE2*∷3xFLAG-E2_K51A_ at 16°C was not the result of suppressor mutations by resequencing the mini-Tn7 construct of the strain grown at 16°C as well as by assessing the growth of the strain in absence of arabinose (i.e. not inducing RhlE2_K51A_ expression). Instead, the swarming motility of the Δ*rhlE2* mutant expressing 3xFLAG-E2_K51A_ was similar to the Δ*rhlE2* mutant (**Fig 7A**). We also used the *lasB’-‘lacZ* fusion as proxy of RhlE2 regulatory activity, as it was downregulated in the Δ*rhlE2* mutant (**Fig 4D**). Again, the expression of the fusion was similarly affected in the Δ*rhlE2* mutant expressing RhlE2_K51A_ as in the Δ*rhlE2* mutant, while the 3FLAGxRhlE2 construct was restoring the expression of *lasB’-‘lacZ* in the Δ*rhlE2* mutant to the wild type levels (**Fig S8**). In all these conditions, unlike at 16°C, the catalytic activity of RhlE2 is essential for its regulatory action.

**Figure 7.**
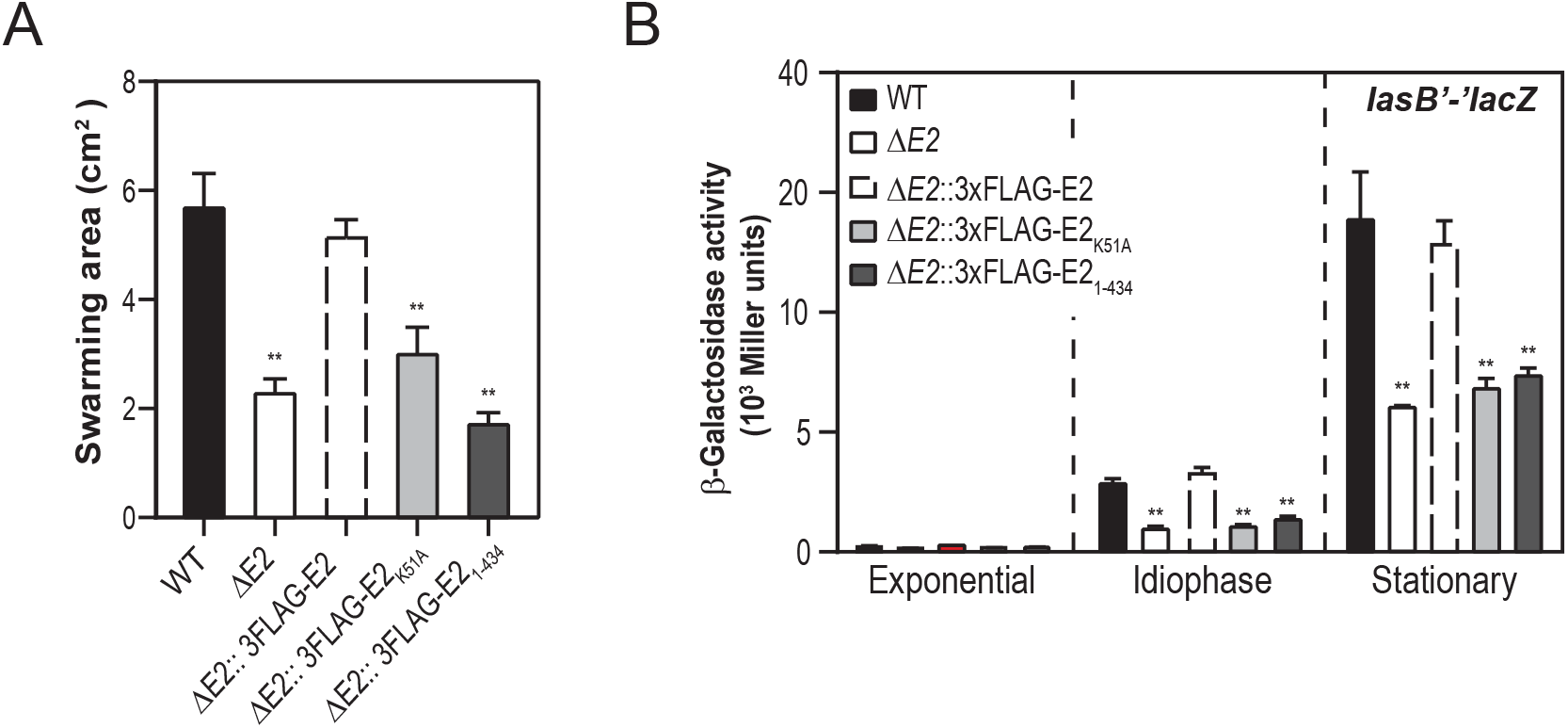
Complementation assays with RhlE2 variants. **(A)** Swarming motility and **(B)** *lasB’-‘lacZ* expression of wild type PAO1, Δ*rhlE2* mutant (ΔE2) and Δ*rhlE2* mutant carrying a mini-Tn7 construct expressing either 3xFLAG-tagged RhlE2 (∷3xFLAG-E2) or 3xFLAG-tagged RhlE2_1-434_ (∷3xFLAG-E2_1-434_), or 3xFLAG-tagged RhlE2_K51A_ (∷3xFLAG-E2_K51A_). Each experiment was repeated at least with three different independent cultures for each strain, values indicated average from biological replicates ± standard deviation. For experimental details see Material and Methods.

### RhlE2 interacts with the Ribonuclease E via RNA

One possibility to explain the different RhlE1 and RhlE2 regulatory actions was that they interact with different protein partners. To test this hypothesis, we probed RhlE1 and RhlE2 proteins interacting partners by protein affinity assays (pull-downs) using *P. aeruginosa* Δ*rhlE1*∷3xFLAG-RhlE1 and the Δ*rhlE2*∷3xFLAG-RhlE2 strains, respectively. Strains were grown at 37°C to an OD600 of ~1.5 in presence of 0.02% of arabinose and the proteins were purified using anti-FLAG antibody resin (see Material and Methods). The protein profile of the eluted fractions was first analysed by SDS page and then by mass-spectrometry (**Fig 8A)**. As control, we used an Δ*rhlE2* strain expressing untagged RhlE2 (CN). Two bands appeared specifically in the elution profile of the 3xFLAG-RhlE2 construct, one at ~80 kDa corresponding to 3xFLAG-RhlE2 and one additional band at ~140 kDa that was excised from the gel and identified by mass-spectrometry as RNase E. On the other hand, no band other rather than RhlE1 was visible in the elution of the 3xFLAG-RhlE1 pull-down (**Fig 8B**). To identify putative partners that were not visible as clear bands in denaturing gels stained with Coomassie, the entire eluted samples were further analysed by mass-spectrometry. Additional proteins were consistently enriched more than 2-fold in the 3xFLAG-RhlE2 elution of three independent pull-down experiments when compared to the negative control (**Fig 8C**). Interestingly, the three most enriched proteins in the 3xFLAG-RhlE2 sample were ribonucleases: RNase E, PNPase and RNase R; suggesting a role of RhlE2 in RNA processing and degradation. The fourth most enriched protein was the DnaK chaperone, which appeared enriched as well in the 3xFLAG-RhlE1 sample.

**Figure 8.**
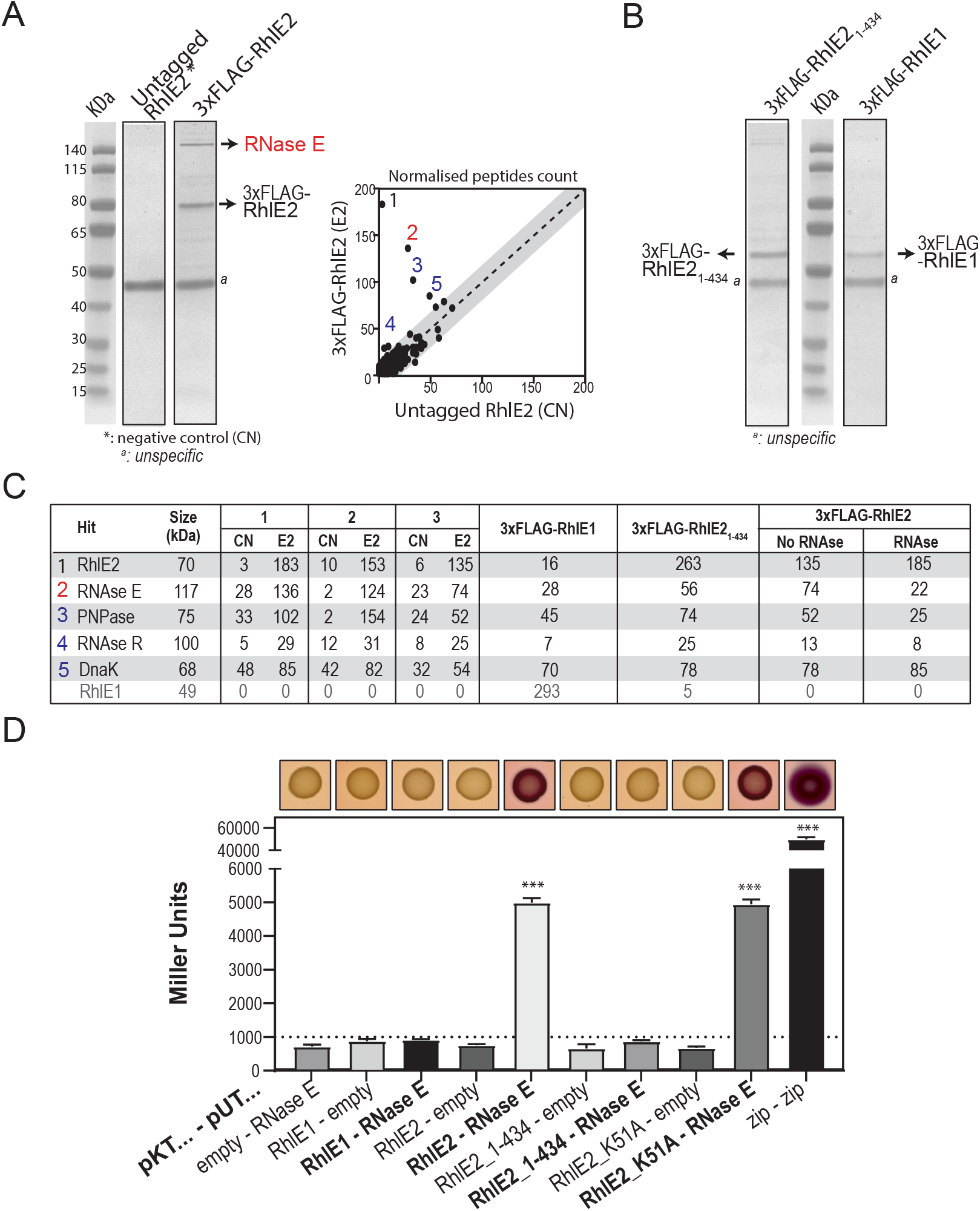
RhlE2 - RNase E interaction. Pull-down assays of 3xFLAG-RhlE2, 3xFLAG-tagged RhlE2_1-434_ and 3xFLAG-RhlE1 and SDS-page analysis of co-eluting proteins. Culture of **(A)** Δ*rhlE2* mutant carrying a mini-Tn7 construct expressing either 3xFLAG-tagged RhlE2 (3xFLAG-RhlE2) or RhlE2 (untagged RhlE2) or of **(B)** Δ*rhlE2* mutant carrying a mini-Tn7 construct expressing 3xFLAG-tagged RhlE2_1-434_ (3xFLAG-RhlE2_1-434_) and Δ*rhlE1* mutant carrying a mini-Tn7 construct expressing 3xFLAG-tagged RhlE1 (3xFLAG-RhlE1) were grown on NYB with 0.05% arabinose at 37°C up to idiophase of growth (O.D. ~1.8). Arrowheads proteins were identified by mass-spectrometry. Other proteins co-eluting with 3xFLAG-RhlE2 were identified by mass-spectrometry and comparison of normalized peptide counts present in the untagged RhlE2 sample, as shown in panel A. **(C)** Quantification of top three proteins which peptides were significantly enriched in 3xFLAG-RhlE2, meaning RNase E, PNPase and RNase R as compared to other strains. Pull-down of 3xFLAG-RhlE2 was performed in triplicates (1, 2 and 3), as well in absence (no RNase) and presence (RNase) of RNase A treatment. **(D)** Reconstitution of adenylate cyclase in the *E. coli* strain BTH101 using a bacterial two-hybrid approach was detected by red colour of colonies due to media acidification derived from maltose fermentation when colonies were grown on McConkey plates containing 1% maltose, 0.5 mM IPTG, 100 μg/ml ampicillin, and 50 μg/ml chloramphenicol agar plates, as shown). The interactions were also quantified in Miller Units by β-galactosidase assays using liquid cultures of the same strains. Each value is the average of three different cultures ± standard deviation (*, *p* < 0.05; **, *p* < 0.01).

Next, we wondered whether the observed RhlE2-protein interactions were direct protein-protein interactions or interactions mediated by RNA. To address this question, additional 3xFLAG-RhlE2 pull-downs were performed by pre-treating cells with RNase A. The enrichment of peptides corresponding to RNase E, PNPase and RNase R was lost in the 3xFLAG-RhlE2 elution sample treated with RNase A, when compared to the untreated sample, attaining background levels comparable to the negative control; whereas peptides corresponding to DnaK were still present (**Fig 8C**). Thus, we conclude that RNA is necessary to mediate or stabilize any interaction of RhlE2 with RNases.

To confirm the pull-down results obtained, we performed bacterial two-hybrid assays, which report on protein interactions based on proximity of split T25-T18 adenylyl cyclase domains (47). Co-expression of T25-*rhlE2* and T18-*rne*, *rne* encoding RNase E, restored the enzyme activity and resulted in red colonies when cells were grown in Congo Red medium and in a significant increase of β-galactosidase activity compared to controls (**Fig 8D**). In agreement with the pull-down assays, T25-RhlE1 interaction with T18-RNase E was not detected. In addition, the K51A mutation did not affect interaction of RhlE2 with T18-RNase E. Finally, interaction of RhlE2 with RNase R and PNPase was not detected via bacterial two-hybrid assays, indicating that these interactions probably occur indirectly via *P. aeruginosa* RNase E (**Fig S9**).

### The CTEs of RhlE2 is necessary for interaction with RNase E and the protein function in vivo

In some cases, the CTE of RNA helicases is necessary for their function in vivo, being involved in mediating interaction with other proteins and/or in recognition of RNA targets (8,48,49). RhlE2 possesses a unique CTE that differs from RhlE1 and from other RhlE homologs investigated so far (**Fig S2**). We asked whether the RhlE2 CTE was important for its biological function by constructing a mini-Tn7 construct carrying 3xFLAG-RhlE2 truncated of its C-terminal region (3xFLAG-RhlE2_1-434_) and performing complementation assays with Δ*rhlE2*∷3xFLAG-RhlE2_1-434_ strain (**Fig S4**). Deletion of the CTE RhlE2 resulted in the inability to restore all the phenotypes tested, *i.e.* growth at 16°C and swarming motility as well as the inability to restore expression of the *lasB’-‘lacZ* reporter fusion to wild type levels (**Fig 2D and Fig 7**).

Then, the RhlE2-RNase E interaction was probed via pull-down assays in a Δ*rhlE2*∷3xFLAG-RhlE2_1-434_ strain and via co-expression of T25-RhlE2_1-434_ and T18-RNase E followed by bacterial two-hybrid assays. With pull-downs, we could show that the interaction with RNase E was lost (compare the level of RNase E on Coomassie gel in **Fig 8A** and in **Fig 8B**). The analysis of the eluates by mass-spectrometry still reveals the presence of some RNase E peptides (56 peptides in 3XFLAG-RhlE2_1-434_ versus 74 peptides in 3XFLAG-RhlE2; **Fig 8C**). However, the results with the bacterial two hybrids showed unambiguously the involvement of the RhlE2 CTE in RNase E binding in vivo (**Fig. 8D**).

Having shown that the unique CTE of RhlE2 was necessary to interact with RNase E, we next asked whether the CTE was necessary for RhlE2 catalytic activity. To address this question, the RhlE2_1-434_ mutant was expressed in *E. coli* with an N-terminal His_10_-Smt3-tag and purified as described previously for RhlE2 (**Fig 9A**). The recombinant RhlE2_1-434_ was still able to catalyze the release of ^32^Pi from [*γ*- ^32^P] ATP in the presence of poly(U) RNA and the extent of ATP hydrolysis was proportional to enzyme concentration, with 56% of the ATP substrate was hydrolyzed in 15 minutes reaction at 100 ng/μl RhlE2_1-434_ (**Fig 9B**). A specific activity of 0.011 nmol of ATP hydrolyzed per ng of protein in 15 min was calculated from the slope of the titration curve in the linear range. This value translates into a turnover number of 46 min^−1^, which is similar to the RhlE2 wild type turnover (51 min^−1^, **Fig 6B**). The observed RNA independent phosphohydrolase activity of RhlE2_1-434_ is intrinsic to the recombinant protein as insofar the ATPase activity of the RhlE2_K51A,1-434_ mutant in the presence poly(U) RNA was less than 1% of the activity of RhlE2_1-434_ (**Fig 9C**). From these results, we conclude that the CTE of RhlE2 (amino-acid 434 to 648) is necessary for the protein regulatory action in vivo and its interaction with RNase E, while it is dispensable for its RNA-dependent ATPase activity in vitro.

**Figure 9.**
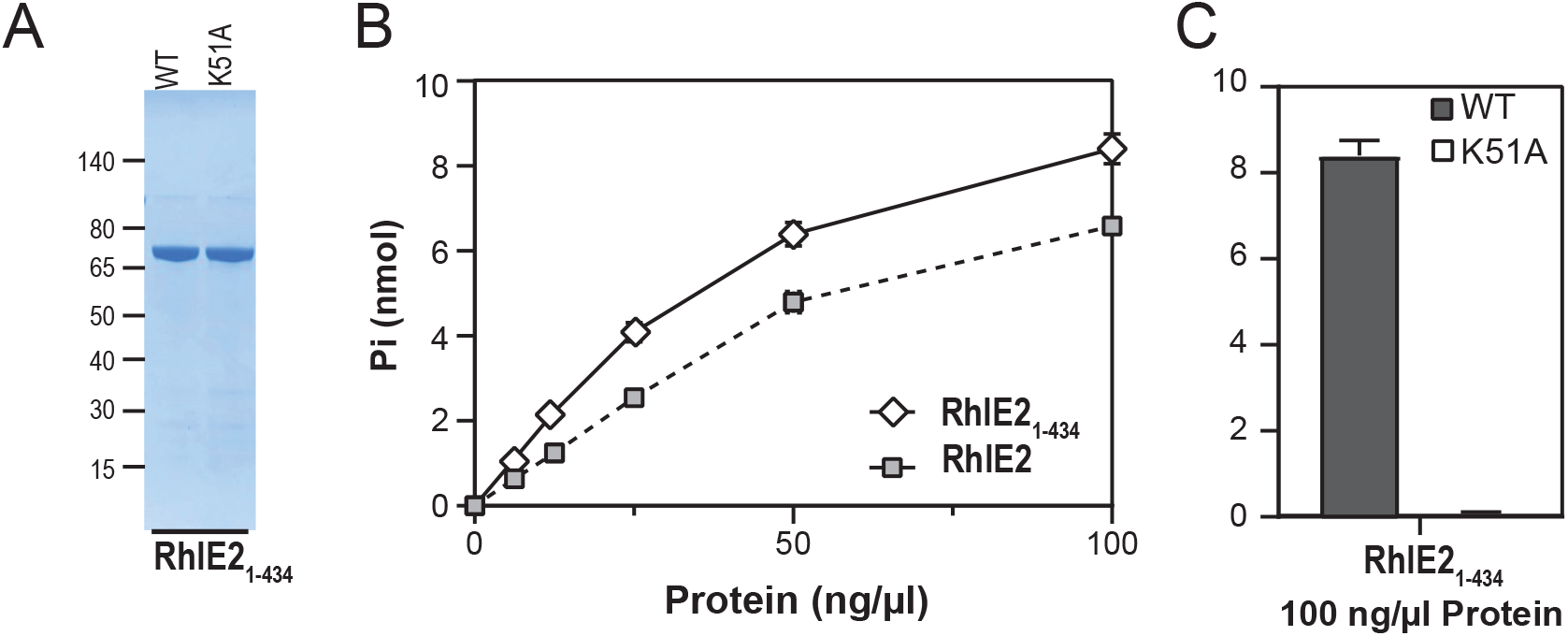
ATPase activity of recombinant RhlE2_1-434_. **(A)** RhlE2_1-434_ purification. Aliquots (2.5 μg) of the nickel-agarose preparations of RhlE2_1-434_ (lane 1) and double mutant RhlE2_1-434_, _K51A_ (lane 2) were analyzed by SDS-PAGE. The polypeptides were visualized by staining with Coomassie Blue dye. The positions and sizes (kDa) of marker polypeptides are indicated on the left. (B) ATPase. Reaction mixtures (15 μl) containing 50 mM Tris-HCl (pH 8.0), 1 mM DTT, 2 mM MgCl2, 1 mM [*γ*-^32^P] ATP, 250 ng/μl Poly(U), and 100 ng/μl of RhlE2 (1–434) or K51A mutant were incubated for 15 min at 37 °C. The extends of ATP hydrolysis are plotted. **(C)** ATPase. Reaction mixtures (15 μl) containing 50 mM Tris-HCl (pH 8.0), 1 mM DTT, 2 mM MgCl2, 1 mM [*γ*-^32^P] ATP, 250 ng/μl Poly(U) and RhlE2(1–434) or K51 mutant protein as specified were incubated for 15 min at 37 °C. Pi release was plotted as a function of input protein. **(B-C)** Data are the average ± SEMs from three independent experiments.

### RhlE2 affects the stability of some cellular RNAs

The RhlE2 interaction with RNase E suggests that RhlE2 could perform a role in cellular RNA decay (50). In order to test this hypothesis, we assessed the half-life of some mRNAs in the wild type and Δ*rhlE2* mutant (**Fig 10**). For the analysis, we selected transcripts whose abundance appeared to be either down-regulated or up-regulated in our RNAseq analysis (**Fig 4**). The mRNA which were up-regulated in the *rhlE2* mutant, like PA1897, PA2314 or PA2268, displayed a 1.5 to 3-fold increase of their half-life compared to the wild type. On the other hand, the stability of *lasB*, a down-regulated mRNAs, do not seem to be affected by *rhlE2* deletion, probably indicating an indirect regulation by RhlE2. In agreement with this hypothesis, we observed a down-regulation of *lasB* expression in the Δ*rhlE2* mutant also when we used a transcriptional *lasB-lacZ* fusion, reporting only on the activity of the *lasB* promoter rather than the mRNA stability or translation rate (**Fig S9**).

**Figure 10.**
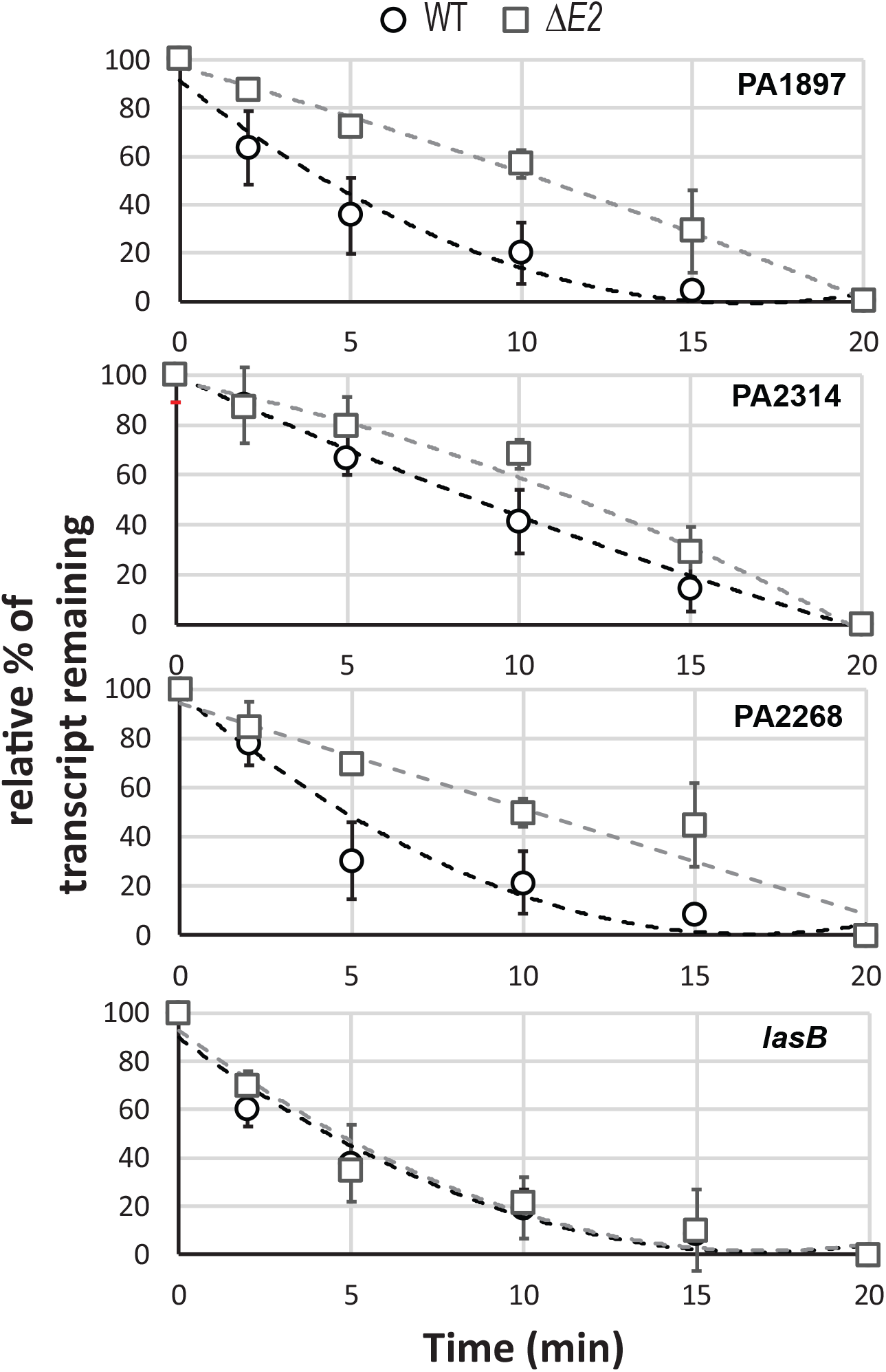
RhlE2 regulation of RNA stability. Determination of the steady state levels and half-life of PA1897, PA2314, PA2268 and *lasB* mRNA by RT-qPCR in the wild type PAO1 and Δ*rhlE2* mutant. Strains were grown in NYB medium until they reached an OD_600_ of 1.5. Then 250 μg/ml rifampicin was added and samples after 1, 5, 10, 15 and 20 minutes were collected for RNA extraction and cDNA synthesis. The steady state levels of mRNAs were normalized to *rpoD* and *rplA* mRNA levels. The level of each mRNA levels at t_0_ was set to 100% while at t_20_ was set to 0 and relative % of transcript remaining is plotted over time. The results represent the average of two independent experiments including each three technical replicates. The error bars represent SDs.

## Discussion

The molecular mechanisms leading to *P. aeruginosa* pathogenesis have been a blooming subject of investigation over the past decades. Numerous toxins, virulence and virulence-associated factors have been shown to be responsible for the variety of infections associated to the bacterium (51). Yet new regulatory networks are still to find. In this paper, we report for the first time the role of a RhlE protein in the regulation of *P. aeruginosa* pathogenesis.

This study was initially motivated by two observations based on a phylogenetic analysis of the RhlE group of proteobacterial RNA helicases. First, genes encoding RhlE proteins have expanded in Proteobacteria through duplication events, reaching up to six copies in *Shewanella pealeana* ATCC 700345 (12), redundancy and sub-functionalization among RhlE proteins had not been investigated. Second, RhlE homologs possess different and fast evolving C-terminal extensions. Whether variability confers a different function to RhlE proteins is not known.

RhlE proteins have been experimentally investigated only in a handful of bacterial species, namely *E. coli*, *Y. pseudotuberculosis*, *M. tuberculosis*, *P. syringae* LZ4W (psychotropic strain) and *C. crescentus* (13–19). Importantly, all the bacterial species in which RhlE proteins were studied so far possess one copy of the *rhlE* gene. *P. aeruginosa* genome owns two copies; therefore, it represents an ideal model organism to investigate gene evolution following duplication, assessing conserved and new gene functions. Our phenotypic analysis of *rhlE1* and *rhlE2* mutants confirmed a widespread role of RhlE proteins in sustaining cell growth at cold temperatures, as both *P. aeruginosa rhlE1* and *rhlE2* mutants display a growth defect at 16°C and a growth profile similar to the wild type at 37°C. Cold adaptation is one of the most frequent biological functions associated with RNA helicases. The function is linked to ability of RNA helicases to resolve RNA secondary structures, which are stabilized at low temperatures, allowing the expression of gene(s) essential for growth on cold (15).

In addition to cold adaptation, deletion of *rhlE2* in *P. aeruginosa* affects cell motility, reduces biofilm formation and production of virulence factors, ultimately resulting into a significant decrease of lethality in *G. mellonella* infection model. Few other groups of DEAD-box RNA helicases, namely CshA of *Listeria monocytogenes* and *Staphylococcus aureus* and RhpA of *Helicobacter pylori* have been shown to regulate expression of virulence traits (52–54). In addition, another *P. aeruginosa* RNA helicase (PA2840, named DeaD) has been proposed as virulence regulator, due to its stimulatory effect on the synthesis of the type III secretion system regulator ExsA (30). Nonetheless, such an important action as the one of RhlE2 on in vivo host-pathogen interaction is rarely described for a DEAD-box RNA helicase and it has never been observed before for RhlE proteins. Phylogenetic analysis of RhlE proteins indicate that RhlE2 homologs are present in other bacteria phylogenetic distant from *P. aeruginosa*, including environmental bacteria like *Azotobacter vinelandii* and several other bacterial pathogens like *Klebsiella pneumoniae*, *Acinetobacter baumannii*, *Enterobacter clocae* and *Streptococcus dysgalactiae* subsp. *equisimilis* (**Fig S1**), suggesting gene acquisition by horizontal gene transfer. It would be interesting to know whether RhlE2 also plays a role on virulence in other pathogenic species.

Contrary to *rhlE2*, *rhlE1* mutation doesn’t produce a pleiotropic effect and, when expressed *in trans*, the RhlE1 protein is unable to complement the *rhlE2* mutant phenotypes, indicating that the two proteins act on different targets. In vitro, RhlE1 and RhlE2 possess an ATPase activity strictly RNA-dependent. No base specificity has been observed in our ATPase assays (see 41-mer ssRNA vs U41), but this is not surprising since the RNA helicase catalytic core usually interacts with the 2’ OH ribose or the phosphate of the RNA backbone (55). Instead, different RNA preferences are observed depending on RNA length and structure: RhlE1 ATPase is activated by shorter RNA than RhlE2. However, RhlE1 is not sensitive to the RNA structure whereas RhlE2 prefers RNA with a hairpin than RhlE2 (**Fig. 6E**). The difference in RNA length stimulation can be explained by a larger RNA-binding site than on RhlE1 or a cooperative binding to RNA. The structure specificity may reflect a hairpin recognition domain on the C-terminal extension. An in-depth biochemical characterization is needed to explore how RhlE2 can discriminate RNA secondary structure.

At the molecular level, the *P. aeruginosa rhlE2* deleted strain present changes in 15% of the entire cellular transcripts in swarming conditions (N=836 transcripts). Identifying direct targets and assessing interconnection with other regulatory pathways will be the subject of further inquiry. Surprisingly, no discernible changes were observed in the mRNA levels of flagellar and rhamnolipids biosynthesis genes, suggesting that the observed swarming defect of the *rhlE2* mutant might result from a metabolic or energetic imbalance. Another possibility is that the reduced swarming motility of the strain might be linked to the strong down-regulation of the virulence factors elastase LasB and PvdQ, which production is interestingly required for swarming motility (33).

In our work, we also show that the RNase E endonuclease, key degradosome component, is co-eluted with FLAG-tagged RhlE2, but not with RhlE1. While studying the giant phage ϕKZ gp37/Dip protein, which interacts with RNase E, Van den Bossche and colleagues recently identified RhlE2 as a possible component of the *P. aeruginosa* degradosome, but the protein was called DeaD as for DEAD-box protein (56). Interaction of RhlE proteins with RNase E is conserved across several bacterial species as it has also been observed previously in *E. coli*, *C. crescentus*, *P. syringae* LZ4W and *M. tuberculosis* (13,17,18,20). In *E. coli*, the RhlE interaction with RNAse E has been observed only in vitro, where the protein can replace the RNA helicases RhlB in facilitating the degradation of structured RNA by PNPase (20). *E. coli* RhlB remains the only helicase with clear role in RNA degradation in vivo and the physiological conditions in which RhlE can replace RhlB are still unknown (57). A RhlB homolog is also present in *P. aeruginosa*, but deletion of the encoding gene doesn’t lead to the same phenotypes observed for *rhlE2* mutant (data not shown). Surprisingly, pre-treatment of *P. aeruginosa* cell extracts with RNase A before pull-down affects RhlE2 interaction with RNase E. The same experiment was performed in *C. crescentus*, but in that case the interaction resisted to the treatment (13). These results indicate that, while in *C. crescentus* interaction of RhlE with RNase E is a protein-protein interaction, *P. aeruginosa* interaction of RhlE2 with RNase E is mediated or stabilized by RNA. The C-terminal extension of RhlE2 is necessary for the observed interaction, as RNase E does not interact with RhlE2_1-434_ construct in bacterial two-hybrid assays (**Fig S8**). The RhlE2_1-434_ protein shows in vitro an RNA-dependent ATPase activity similar to the full-length protein, but it does not restore in vivo the *rhlE2* mutant phenotypes to wild type levels, suggesting that RhlE2 performs its regulatory action via the RNA degradosome. Indeed, we show that RhlE2 affects the half-life of some of the transcripts whose levels were affected by *rhlE2* deletion in the RNA-sequencing analysis. In *E. coli*, RNase E was shown to interact with RhlB by binding to the RecA2 domain and stimulating its ATPase activity (58,59). We do not observe stimulation of RhlE2 activity by RNase E (data not shown), probably due to the different type of interactions these two helicases enact with the endonuclease. Our work on RhlE2 hints that *P. aeruginosa* RNA degradosome mode of action might differ from *E. coli*, as also suggested when looking at RNase E sequence conservation among these two species (60). We are currently investigating this matter.

## Acknowledgements

We are grateful to Patrick Linder for critical reading of the manuscript. RNA-sequencing experiments were performed at the iGE3 genomics platform of the University of Geneva while MS/MS protein identifications were performed at the proteomics core facility of the Faculty of Medicine (University of Geneva), and we are very grateful for their help and advices.

## Materials and Methods

### Bacterial strains and culture conditions

Bacterial strains and plasmids used in this study are listed in **Table S4**. Cells were grown in Luria Broth (LB) or nutrient yeast broth (NYB) (61,62) medium, with shaking at 180 rpm and at 37°C. Nutrient agar (NA) was used as a solid medium. When required, antibiotics were added to these media at the following concentrations: 100 μg/ml ampicillin, 25 μg/ml tetracycline and 10 μg/ml gentamicin for *E. coli*; and 50 μg/ml gentamicin and 50 μg/ml tetracycline for *P. aeruginosa*.

### Genetic techniques

DNA cloning and plasmid preparation were performed according to standard methods (63). Oligonucleotides used for cloning procedures are listed in **Table S5**. Restriction and DNA-modifying enzymes were used following the instructions of the manufacturers. Transformation of *E. coli* DH5α, *E. coli* TOP10 (for cloning) and *P. aeruginosa* was carried out by heat-shock and electroporation, respectively (64). The construction of engineered plasmids and strains is detailed in Supplemental Material. All plasmids and strains were verified by PCR and sequencing.

### RhlE1 and RhlE2 purifications

The pET28-His_10_Smt3-RhlE1 and RhlE2 plasmids were transformed into *E. coli* BL21(DE3). Cultures (500 ml) derived from single transformants were grown at 37°C in LB medium containing 50 μg/ml kanamycin until the OD_600_ reached 0.6. The cultures were adjusted to 0.2 mM IPTG and 2% (v/v) ethanol and incubation was continued for 20 h at 17°C. Cells were harvested by centrifugation and stored at −80°C. All subsequent procedures were performed at 4°C. Thawed bacteria were resuspended in 25 ml of buffer A (50 mM Tris-HCl, pH 8.0, 500 mM NaCl, 10% glycerol) and supplemented with one tablet of protease inhibitor cocktail (Roche). The suspension was adjusted to 0.1 mg/ml lysozyme and incubated on ice for 30 min. Imidazole and Triton-X100 were added to a final concentration of 10 mM and 0.1%, respectively. The lysate was sonicated to reduce viscosity and insoluble material was removed by centrifugation. The soluble extracts were applied to a 1 ml of Ni^2+^- NTA-agarose (Qiagen) column that had been equilibrated with buffer A containing 10 mM imidazole. The column was washed with 10 ml aliquots of 20 mM imidazole in buffer A and then eluted stepwise with 3 ml aliquots of buffer A containing 50, 100, 250, and 500 mM imidazole, respectively. The elution profiles were monitored by SDS-PAGE (not shown; 250 mM imidazole fraction is shown in Fig 5A).

The recombinant His10Smt3-RhlE polypeptides were recovered predominantly in the 250 mM imidazole fractions. The protein concentration of the 250 mM imidazole eluate fraction was determined using the Bio-Rad dye reagent with BSA as the standard. Proteins concentration were calculated by interpolation to the BSA standard curve. RhlE1 and RhlE2 mutated variants were produced and purified as described above for the wild type RhlE1 and RhlE2.

### ATPase reaction

Reaction mixtures (15 μl) containing 50 mM Tris-HCl, pH 8.0, 5 mM DTT, 2m MgCl2, 1mM ATP, RNA as specified, and enzyme as specified were incubated for 15 min at 37°C. The reactions were quenched by adding 3.8 μl of 5 M formic acid. Aliquots (2 μl) were applied to a polyethylenimine(PEI)-cellulose thin layer chromatography (TLC) plates (Merk), which were developed with 1 M formic acid, 0.5 M LiCl. ^32^Pi release was quantitated by scanning the chromatogram with laser Scanner Typhoon FLA 7000 (General Electric).

### β-Galactosidase assays

β-Galactosidase experiments were performed as described previously (62), with *P. aeruginosa* strains grown in NYB medium. Data are mean values of three independent samples ± standard deviations.

### Phenotypic assays

*Growth assays.*Liquid growth assays at 25, 37 and 43°C were performed in 96-well plates each containing 200 μl of LB and cell density (OD 600) was measured with a Synergy H1 plate reader (Biotek). Spot assays were performed by spotting 10 μl of serial dilutions of cultures (at initial OD600 of 1.0) on LB plates incubated at 16 °C or at 37 °C. *Biofilm formation.* Quantification of biofilm formation was performed in 24-well polystyrene microtiter plates as previously described (65). The plates were incubated for 10 h at 37°C and biofilms were stained with 0.1% crystal violet solution. The dye bound, which is proportional to the biofilm produced, was solubilized with 96% (v/v) ethanol and the absorption was photometrically measured at 600 nm (OD_600_). Data are mean values of three independent samples ± standard deviations. *Motility assays.* Motility assays were carried out essentially as previously described (66,67). Swim assays were carried out on 10 g/L tryptone, 5 g/L NaCl, 0.3% agar (Merck) plates. 0.5 μl of standardized overnight culture was injected below the surface of the agar and plates were incubated at 30°C overnight. Twitch assays were carried out on 1% LB agar plates and bacteria were inoculated by picking a colony using a sterile tip and stabbing to the bottom of the plates, which were incubated at 37°C for two days. The agar was then peeled off the plate, and cells were stained with crystal violet for visualization. Swarming assays were performed in plates consisted of 0.5% (wt/vol) Difco bacto-agar with either 8 g/liter Difco nutrient broth and 5 g/liter glucose or with MMP MMP medium supplemented with 20 mM glucose and 0.1% (w/v) casamino acids (66,68). Cells were inoculated onto swarm plates using 5μl of standardized overnight cultures and incubated overnight at 37°C. Pictures are taken from a representative plate out of five independent experiments. Swarming complementation experiments were performed in 0.5% (wt/vol) nutrient agar plus 0.2% arabinose. Pictures are taken from a representative motility plate out of five independent experiments. ImageJ software (NIH) was used to determine the area of the plate surface covered by the bacteria as previously described (69).

### Bacterial two-hybrid assay

Bacterial two hybrid experiments (70). Recombinant pKT25 and pUT18C derivative plasmids were transformed simultaneously into the *E. coli* BTH101 strain. Transformants were spotted onto McConkey agar plates supplemented with 1mM isopropyl β-d-thiogalactoside (IPTG) in the presence of 100 μg/ml ampicillin, 50 μg/ml kanamycin, and 100 μg/ml 5-bromo-4-chloro-indolyl-β-d-galactopyranoside (X-gal). Positive interactions were identified as blue colonies after 24h incubation at 30°C and quantified by β-galactosidase assays. The positive controls used in the study were pUT18C and pKT25 derivatives encoding the leucine zipper from GCN4 (zip), which strongly dimerizes. Data are mean values of three independent samples ± standard deviations.

### Virulence factor production assays

*Elastase production.* The LasB elastase activity of bacterial suspensions was determined with the Elastin-Congo red (ECR; Sigma, St. Louis, MO) assay, as previously described (71). Briefly, 50 μl aliquots of filtered bacterial supernatant of a 21 h culture were added to 900 μl of ECR buffer (100 mM Tris, 1 mM CaCl2, pH 7.5) containing 20 mg of ECR and then incubated with shaking at 37°C for 18 h. Insoluble ECR was removed by centrifugation and the absorption of the supernatant was measured at 495 nm and normalized according to cell density (OD600). LB medium was used as a negative control*. Pyocyanin production.* Pyocyanin quantification was performed using the assay based on absorbance at 520 nm in an acidic solution. Briefly, 5 ml supernatant from a stationary phase culture was mixed with 3 ml of chloroform. Pyocyanin from the organic phase was then extracted with 1 ml of 0.2 N HCl, and absorbance was measured at 520 nm. Concentrations, expressed as micrograms of pyocyanin produced per milliliter of culture supernatant, were determined by multiplying the optical density at 520 nm by 17.072 (72). The experiment was conducted three times in independent experiments.

### In vivo virulence assay

The wax moth model *Galleria mellonella* was used for *in vivo* virulence assays, as previously described (73). Briefly, overnight cultures were diluted 1:100 and sub-cultured (3 ml LB) at 37°C until exponential phase (3 hours). Bacterial cells (OD600 = 1.0) were washed with PBS three times. The samples were serially diluted (up to 10−7) and a 10 μl volume of the dilutions was spotted on LB agar and CFU counted the next day. A 10 μl volume of the 10−7 dilution for the wild type and mutant strains was injected into the last abdominal proleg of ten larvae per sample. An additional ten larvae were injected with PBS as negative control. Larvae mortality was monitored every two hours from 16 to 40 hrs post-infection (larvae are considered dead when they turn black and do not respond to tapping). All data are mean values of three independent samples ± standard deviation.

### Total RNA extraction

PAO1 wild type, PAO1Δ*rhlE1*, PAO1Δ*rhlE2* and PAO1Δ*rhlE1*Δ*rhlE2* mutants were grown at 37°C in swarming condition or on NYB until OD~1.5 with vigorous shaking. Cells were harvested using the RNA bacteria protect solution (QIAGEN). Total RNAs was extracted and purified using Monarch RNA isolation kit (NEB), treated with DNase I (Promega) three times to remove contaminating genomic DNA and re-purified again using phenol-chloroform. Eventual DNA contamination was tested by PCR with 40 cycles and different couples of primers (**Table S5**) and RNA integrity was controlled by agarose gel electrophoresis.

### RNA sequencing

PAO1 wild type, PAO1Δ*rhlE1*, PAO1Δ*rhlE2* and PAO1Δ*rhlE1*Δ*rhlE2* mutants were grown in twelve 12-cm square swarming plates. After O/N incubation at 37°C, samples from 6 plates were pooled and total RNA was isolated from as described above. Two replicates of either strain were therefore obtained. Ribosomal RNA was depleted with Ribo-Zero rRNA Removal Kit (Illumina). Then, libraries were prepared using the Illumina TruSeq stranded mRNA kit and validated on the Bioanalyzer 2100 (Agilent). Samples were sequenced using the Illumina HiSeq 2000, 100 bp single end read at the iGE3 genomics platform of the University of Geneva. After removal of the adaptors, the sequences were quality trimmed with trimmomatic using default parameters (74). The resulting sequences were then mapped onto the PAO1 reference genome (NC_002516.2) using Segemehl (74). For differential gene expression analysis, reads were counted using the DESeq2 package (75). All RNAs with a log2-fold change greater than 2 and a multiple testing adjusted p-value below 0.05 were considered differentially abundant.

### Real-time quantitative PCR (RT-qPCR)

cDNAs from RNA isolated sample was obtained using M-MLV reverse transcriptase (Promega) following the manufacturer instructions. Resultant cDNAs were used as template for qPCR and qPCR was performed on an Applied Biosystems 7900HT Sequence Detection system using a Sybr Green Quantitect kit (QIAGEN, Hildesheim, Germany). Quantification cycle (Cq; standard name for Ct or Cp value) values were recorded with SDS version 2.3 software. Cq values ≥36 were considered beyond the limit of detection. Fold changes were determined using the relative quantification comparative threshold cycle (C_t_) method (ΔΔC_t_) and normalization was done by comparison with *rplA*, *oprD* and *rpoD* housekeeping genes as reference genes. The primer pairs that were used for qRT-PCR are shown in **Table S3**. For each gene tested, a standard curve was made using serial dilutions from 50 to 0.005 ng of *P. aeruginosa* PAO1 genomic DNA in order to quantify the efficiency of every PCR reaction. The PCR products were ranging between 150 and 200 bp in length. All data are mean values of three independent samples and three technical replicates.

### RNA decay analysis

For decay analysis, rifampicin to 250 μg/ml was added to PAO1 wild type, PAO1Δ*rhlE2* liquid culture grown to OD600 of ~1.5 and aliquots were withdrawn at the indicated times. RNA was extracted and used for RT-qPCR analysis using primers listed in **Table S3**.

### Pull-down assays

PAO1Δ*rhlE1*∷P_BAD_-3xFLAG_RhlE1, PAO1Δ*rhlE2*∷P_BAD__RhlE2, PAO1Δ*rhlE2*∷P_BAD_-3xFLAG_RhlE2 and PAO1Δ*rhlE2*∷P_BAD_-3xFLAG_RhlE2_1-434_ cultures were grown in 2L flasks with 400 ml NYB plus 0.05% arabinose to an O.D ~1.8-2.0, then pelleted by centrifugation and stored at −80°C for at least 12 hours. Pellets were re-suspended in ice-cold IP buffer (20 mM HEPES pH 7.4, 300 mM NaCl, 1 mM EDTA, protease inhibitor) and cells were lysed by sonication. Samples were then centrifuged (15,000 g, 60 min, 4°C), the supernatant (soluble fraction) was collected and 1.0% v/v Triton X-100 was added and incubated at 4°C with end-over-end agitation for 20 minutes. Then, samples were incubated for 4 hours with 20 μg/ml ANTI-FLAG M2 affinity gel (Sigma) (4°C, end-over-end agitation). Samples were pelleted by centrifugation (3,000 g, 1 min 4°C), the supernatant was discarded, and the beads re-suspended in 1.0 ml ice-cold IP buffer. This wash step was repeated 5 times. Elution was performed twice in 100 μl 3xFLAG peptide (Sigma) at final concentration of 100 μg/ml. The presence of Flag-tagged RhlE proteins was confirmed by immunoblotting, and co-eluting proteins were detected by Orbitrap mass spectrometry or by SDS-page followed by protein band extraction and identification by mass spectrometry.

### Phylogenetic analysis

Sequences were retrieved from the NCBI RefSeq database (76) and aligned with MUSCLE (77). Sequences of RecQ (PA3344) were added as outgroup.

