## Supplementary material for "The DEAD-box RNA helicases RhlE2 is a global regulator of *Pseudomonas aeruginosa* lifestyle and pathogenesis" (from another version of this preprint: version 1, 2021-01-29)

**Supplemental information**

### Supplementary methods

**Construction of plasmids.** Below description of construction of relevant plasmids is described.

Plasmid pLM41 (pJN_3xFLAG-RhlE2) was constructed by inserting a 1920-bp - fragment, carrying the 3xFLAG-*rhlE2* gene, into pJN105 previously digested EcoRI-SpeI. The 1920-bp fragment was obtained by PCR using the EcoRI-3XFLAG-tagged primer pJN-M2-RhlE2.fw and the SpeI-tagged primer pJN-RhlE2.rev to amplify the PAO1 genome carrying the *rhlE2* gene.

Plasmid pLM88 (pME6182_$P_{BAD}$-3xFLAG_*rhlE2*) was achieved by recruiting an *araC*-$P_{BAD}$-3xFLAG-*rhlE2* fragment from the plasmid pJN-3xFLAG-RhlE2 by PCR with primers pTn7-3F-RhlE2.fw and pTn7-3F-RhlE2.rev; and cloned into the mini-Tn*7* vector pME6182 by Gibson assembly® (NEB).

Plasmid pLM42 (pJN_RhlE2) was achieved by amplification of *rhlE2* by PCR with EcoRI-tagged primer pJN-RhlE2.fw and SpeI-tagged primer pJN-RhlE2.rev and cloning into pJN105 digested EcoRI-SpeI.

Plasmid pLM89 (pME6182_$P_{BAD}$-*rhlE2*) was achieved by recruiting an *araC*-$P_{BAD}$-3xFLAG-*rhlE2* fragment from the plasmid pJN-RhlE2 by PCR with primers pTn7-3F-RhlE2.fw and pTn7-3F-RhlE2.rev; and cloned into the mini-Tn*7* vector pME6182 by Gibson assembly® (NEB).

Plasmid pLM128 (pME6182_$P_{BAD}$-3xFLAG_*rhlE1*) was achieved by amplifying the PAO1 genome carrying the *rhlE1* gene by PCR with primers pTn7-3F-RhlE1.fw and pTn7-3F-RhlE1.rev and cloning into the mini-Tn*7* vector pME6182 by Gibson assembly® (NEB).

For production of His10-Smt3_RhlE1 and His10-Smt3_RhlE2 in *E. coli* BL21 (DE3), the *rhlE1* and *rhlE2* genes were amplified from *P. aeruginosa* genomic DNA (using HindIII-tagged primer pPA3950.fw and XhoI-tagged primer pPA3950.rev or HindIII-tagged primer pPA0428.fw and XhoI-tagged primer pPA0428.rev, respectively) and cloned into pET28b10xHis-Smt3 digested with HindIII-XhoI. The resulting recombinant plasmids pLM38 and pLM37 encoded RhlE1 and RhlE2, respectively, with a N-terminal cleavable 10xHis-Smt3 tag.

For bacterial two-hybrid assays, the *rhlE1* and *rhlE2* genes were amplified from *P. aeruginosa* genomic DNA (using BamHI-tagged primer pBTH_E1.fw and EcoRI-tagged primer pBTH-E1.rev or BamHI-tagged primer pBTH_E2.fw and EcoRI-tagged primer pBTH-E2.rev, respectively) and cloned into pKT25 plasmid digested with EcoRI-BamHI, resulting in pLM86 and pLM87. Moreover, the *rne* gene, encoding RNAse E, was amplified from *P. aeruginosa* genomic DNA (using BamHI-tagged primer pBTH_rne0.fw and EcoRI-tagged primer pBTH_rne4.rev) and cloned into pUT18C plasmid digested with EcoRI-BamHI, resulting in pLM147.

The K51A and ΔCter mutations in the *rhlE1* and *rhlE2* coding sequence were introduced into pLM128 or pLM38 or pLM87 and pLM88 or pLM37 or pLM86, respectively, by PCR. The following mutagenesis primer pairs were used pRhlE1_K51A.fw-pRhlE1_K51A.rev ($RhlE1_{K51A}$), pRhlE1_1-374.fw-pRhlE1_1-374.rev ($RhlE1_{1-374}$), pRhlE2_K51A.fw-pRhlE2_K51A.rev ($RhlE2_{K51A}$), pRhlE2_1-424.fw-pRhlE2_1-434.rev ($RhlE2_{1-434}$). The parental DNA template was digested with DpnI and the mutated plasmid was transformed into *E. coli* DH5α, generating pLM46, pLM47, pLM79, pLM90, pLM91, pLM111, pLM112 and pLM143.

**Construction and description of relevant *P. aeruginosa* strains**. For the inactivation of the *rhlE1* and *rhlE2* gene in the *P. aeruginosa* PAO1 chromosome previously described protocol was followed (1). Briefly, a fragment containing the upstream region (~500-bp) overlapping the first 2 codons of the gene and a ~500-bp downstream fragment overlapping the stop codon were amplified by PCR using the primer couples pPA3950.1/pPA3950.2 and pPA3950.3/pPA3950.4 for *rhlE1* and pPA0428.1/pPA0428.2, and pPA0428.3/pPA0428.4 for *rhlE2*. The upstream and downstream overlapping products were digested with EcoRI-HindIII and cloned into pME3087, giving plasmid pLM2 and pLM3, respectively. Plasmid pLM2 or pLM3 were then introduced into *P. aeruginosa* PAO1 as described by triparental mating, using the helper strain *E.coli* HB101 (pRK2013). Transconjugants were isolated on nutrient agar supplemented with appropriate antibiotics. Merodiploids were resolved as previously described (1) and deletions were confirmed by PCR and sequencing.

The double Δ*rhlE1*Δ*rhlE2* mutant was obtained by crossing plasmid pLM2 into PAO1Δ*rhlE2* (MP4), giving strain MP6 (PAO1Δ*rhlE1*Δ*rhlE2*).

Chromosomal insertion of the pLM88, pLM128 and all mutated variants were introduced into recipient strains by co-electroporation with pUX-BF13, allowing transposition of the constructs into the chromosomal Tn*7* attachment site (2,3).

**Table S1. Strains and plasmids used in this study.**

| **Strains** | **Genotype/relevant characteristics** | **Source** |
|---|---|---|
| *E.coli* | | |
| *Rosetta* (DE3) | *F- ompT hsdSB(rB- mB-) gal dcm* (DE3) pRARE ($Cm^R$) | Novagen |
| DH5α | *recA1 endA1 hsdR17 supE44 thi-1 gyrA96 relA1* Δ(*lacZYA-argF*)*U169* [Φ80d*lacZ*M*15*]$F^-$ $NaI^r$ | (4) |
| HB101 | *proA2 hsdS20*($r_B^-$ $m_B^-$) *recA13 ara-14 lacYI galK2 rpsL20 supE44 xyl-5 mtl-1* $F^-$ | (4) |
| *P. aeruginosa* | | |
| PAO1 | Wild-type | (5) |
| MP3 | PAO1 containing a 1332-bp deletion in the *rhlE1* (PA3950) locus | This study |
| MP4 | PAO1 containing a 1902-bp deletion in the *rhlE2* (PA0428) locus | This study |
| MP6 | PAO1 with a *rhlE1* and *rhlE2* deletion | This study |
| **Plasmids** | | |
| pET28b-10xHis-Smt3 | Broad host range vector for expression of N-terminal 10xHis-Smt3-tag proteins, $Km^r$ | (6) |
| pKT25 | Two-hybrid plasmid, *cyaAT25* fusion, $Km^R$ | (7) |
| pUT18C | Two-hybrid plasmid, *cyaAT18* fusion, $Ap^R$ | (7) |
| pME3087 | Suicide vector for allelic replacement; $Tc^r$; ColE1 replicon | (8) |
| pLM2 | pME3087 derivative for the deletion of *rhlE2* gene, $Tc^r$ | This study |
| pLM3 | pME3087 derivative for the deletion of *rhlE1* gene, $Tc^r$ | This study |
| pME6182 | Mini-Tn*7* gene delivery, ColE1 replicon; $Gm^rAp^r$ | (2) |
| pLM37 | pET28b-10xHis-Smt3 derivative for His-Smt3_RhlE2 purification, $Km^r$ | This study |
| pLM38 | pET28b-10xHis-Smt3 derivative for His-Smt3_RhlE1 purification, $Km^r$ | This study |
| pLM41 | pJN105 derivative for 3xFLAG-RhlE2 expression, $Gm^r$ | This study |
| pLM42 | pJN105 derivative for untagged RhlE2 expression, $Gm^r$ | This study |
| pLM46 | pLM37 derivative with for His-Smt3_$RhlE2_{K51A}$ purification, $Km^r$ | This study |
| pLM47 | pLM37 derivative with for His-Smt3_$RhlE2_{1\text{-}434}$ purification, $Km^r$ | This study |
| pLM79 | pLM38 derivative with for His-Smt3_$RhlE1_{K51A}$ purification, $Km^r$ | This study |
| pLM86 | pKT25 derivative for T25-RhlE2 expression, $Km^r$ | This study |
| pLM87 | pKT25 derivative for T25-RhlE1 expression, $Km^r$ | This study |
| pLM88 | pME6182 derivative carrying $P_{BAD}$_3XFLAG-RhlE2, $Gm^r$ | This study |
| pLM89 | pME6182 derivative carrying $P_{BAD}$_RhlE2 (untagged), $Gm^r$ | This study |
| pLM90 | pME6182 derivative carrying $P_{BAD}$_3XFLAG-$RhlE2_{1\text{-}434}$, $Gm^r$ | This study |

| | | |
|---|---|---|
| pLM91 | pME6182 derivative carrying $P_{BAD}$_3XFLAG-RhlE2$_{K51A}$, Gm$^r$ | This study |
| pLM111 | pME6182 derivative carrying $P_{BAD}$_3XFLAG-RhlE1$_{K51A}$, Gm$^r$ | This study |
| pLM112 | pLM37 derivative with for His-Smt3_RhlE2$_{K51A+1-434}$ purification, Km$^r$ | This study |
| pLM128 | pME6182 derivative carrying $P_{BAD}$_3XFLAG-RhlE1; Gm$^r$ | This study |
| pLM143 | pME6182 derivative carrying $P_{BAD}$_3XFLAG-RhlE1$_{1-374}$; Gm$^r$ | This study |
| pLM147 | pUT18C derivative for T18-RNaseE expression, Ap$^r$ | This study |
| pJN105 | Broad-host-range low-copy-number vector pBBR1-MCS5 harboring araC-PBAD cassette from pBAD18, Gmr | (9) |
| pRK2013 | Helper plasmid; Tra$^+$ Km$^r$ | (4) |

**Table S2. List of cloning primers used in this study.**

| Name | Sequence[a] (5' → 3') | Restriction site[#] |
|---|---|---|
| pPA0428.1 | GGAATTCCTCCGATGCGCCAGTTCAG | EcoRI |
| pPA0428.2 | TTAGCGGTTAAAGGACATGCGGGTTTTCC | |
| pPA0428.3 | ATGTCCTTTAACCGCTAAGGCCGAGCGG | |
| pPA0428.4 | CCCAAGCTTGTGCCAGCCTGGACTACGC | HindIII |
| pPA3950.1 | GGAATTCCCTACGGCTGCCATCTCAAC | EcoRI |
| pPA3950.2 | CTATTTGCCGAACGTCATGAGCGACTCG | |
| pPA3950.3 | ATGACGTTCGGCAAATAGCGAGGCGCTG | |
| pPA3950.4 | CCCAAGCTTGCGAACGGATCGAGGCACG | HindIII |
| PA0428.fw | CAGTAAGCTTCATCCTTTTCTTCCCTCGGACTCTCC | HindIII |
| PA0428.rev | CATGCTCGAGTTAGCGGTTGCGGGTCAGCAG | XhoI |
| pJN_RhlE2.fw | CAGTGAATTCATGTCCTTTTCTTCCCTCGGACTC | EcoRI |
| pJN_M2-RhlE2.fw | CAGTGAATTCATGGACTACAAAGACCATGACGGTGA TTATAAAGATCATGATATCGATTACAAGGATGACGA TGACAAGTCCTTTTCTTCCCTCGGACTCTCC | EcoRI |
| pJN_RhlE2.rev | CATGACTAGTTTAGCGGTTGCGGGTCAGCAG | SpeI |
| pTn7-3F-RhlE2.fw | CGGTTTACAAGCATAAAGCTTCTCGAGGTCGACGGT ATCG | |
| pTn7-3F-RhlE2.rev | GGCAGGCATGCCATGGTACCTCTAGAACTAGTTTAG CGG | |
| pTn7-3F-RhlE1.fw | GATGACGATGACAAGACGTTCGCTTCCCTCGG | |
| pTn7-3F-RhlE1.rev | CAGGCATGCCATGGTACCCTATTTGCCGGACGGCC | |
| pRhlE2_K51A.fw | GACCGGCACCGGCGCGACCGGTGGTTTC | |
| pRhlE2_K51A.rev | GAAACCACCGGTCGCGCCGGTGCCGGTC | |
| pRhlE2_1-424.fw | GGCCCAGGACGGTTGAGAGAAGCCGGCTG | |
| pRhlE2_1-424.rev | CAGCCGGCTTCTCTCAACCGTCCTGGGCC | |

| | | |
|---|---|---|
| pRhlE1_K51A.fw | GACCGGCACCGGCGCGACCGCCGGTTTC | |
| pRhlE1_K51A.rev | GAAACCGGCGGTCGCGCCGGTGCCGGTC | |
| pRhlE1_1-374.fw | GGAGCCCGACTGAGAGCCGGAGCAC | |
| pRhlE1_1-374.rev | GTGCTCCGGCTCTCAGTCGGGCTCC | |
| pPA3950.fw | CAGTAAGCTTCAACGTTCGCTTCCCTCGGTCTGCTC | HindIII |
| pPA3950.rev | CATGCTCGAGCTATTTGCCGGACGGCCGCTTC | XhoI |
| pBTH_E1.fw | GTACGGATCCCACGTTCGCTTCCCTCGGTC | BamHI |
| pBTH-E1.rev | GATCGAATTCCTATTTGCCGGACGGCCGC | EcoRI |
| pBTH_E2.fw | GTACGGATCCCTCCTTTTCTTCCCTCGGACTC | BamHI |
| pBTH-E2.rev | GATCGAATTCTTAGCGGTTGCGGGTCAGC | EcoRI |
| pBTH_rne0.fw | GTACGGATCCCAAAAGAATGCTGATCAACGCGACTC | BamHI |
| pBTH_rne4.rev | GATCGAATTCTCAGGCATGCGGGCGCG | EcoRI |

[#]restriction sites underlined

**Table S3. List of RT-qPCR primers used in this study.**

| Name | Sequence[a] (5' → 3') | Target[#] |
|---|---|---|
| rplA.fw | GCGATTGCCGAGAAAGTAGT | rplA |
| rplA.rev | TCCACGGACTCCTTGAACTT | |
| rhlA.fw | CGGCGCGAAAGTCTGTTGG | rhlA |
| rhlA.rev | GAAATGCCCGGTGCCCTCC | |
| lasB.fw | ACATCGCCCAACTGGTCTAC | lasB |
| lasB.rev | ACTTGCCGATCTTCTGGTTG | |
| pvdF.fw | CGAACAGAAATACTCGGGGC | pvdF |
| pvdF.rev | TTCCTTGGCGATGTCCTTCT | |
| roeA.fw | ATCGCCAGTTGGAGTACCTG | roeA |
| roeA.fw | CCATAGCGGTCGTTGATGTA | |
| lhpF.fw | GTGCTCGTGATGTTCATCGT | lhpF |
| lhpF.rev | AACAGACCGGTGAGTTCGTC | |
| PA2314.fw | GGCCTGTTGTCGCTGACC | PA2314 |
| PA2314.rev | GTGTAGAGGATGTTGTGCGC | |
| PA2268.fw | GCGGCCCGACACCTTATTA | PA2268 |
| PA2268.rev | CAGTAGGAGACGGTGATGCC | |
| PA1897.fw | GCAACGCATCGAGGTATTCA | PA1897 |
| PA1897.rev | CGCAGGTTCGAGTGGTAGAA | |
| oprD.fw | ATCTACCGCACAAACGATGAAGG | oprD |
| oprD.rev | GCCGAAGCCGATATAATCAAACG | |

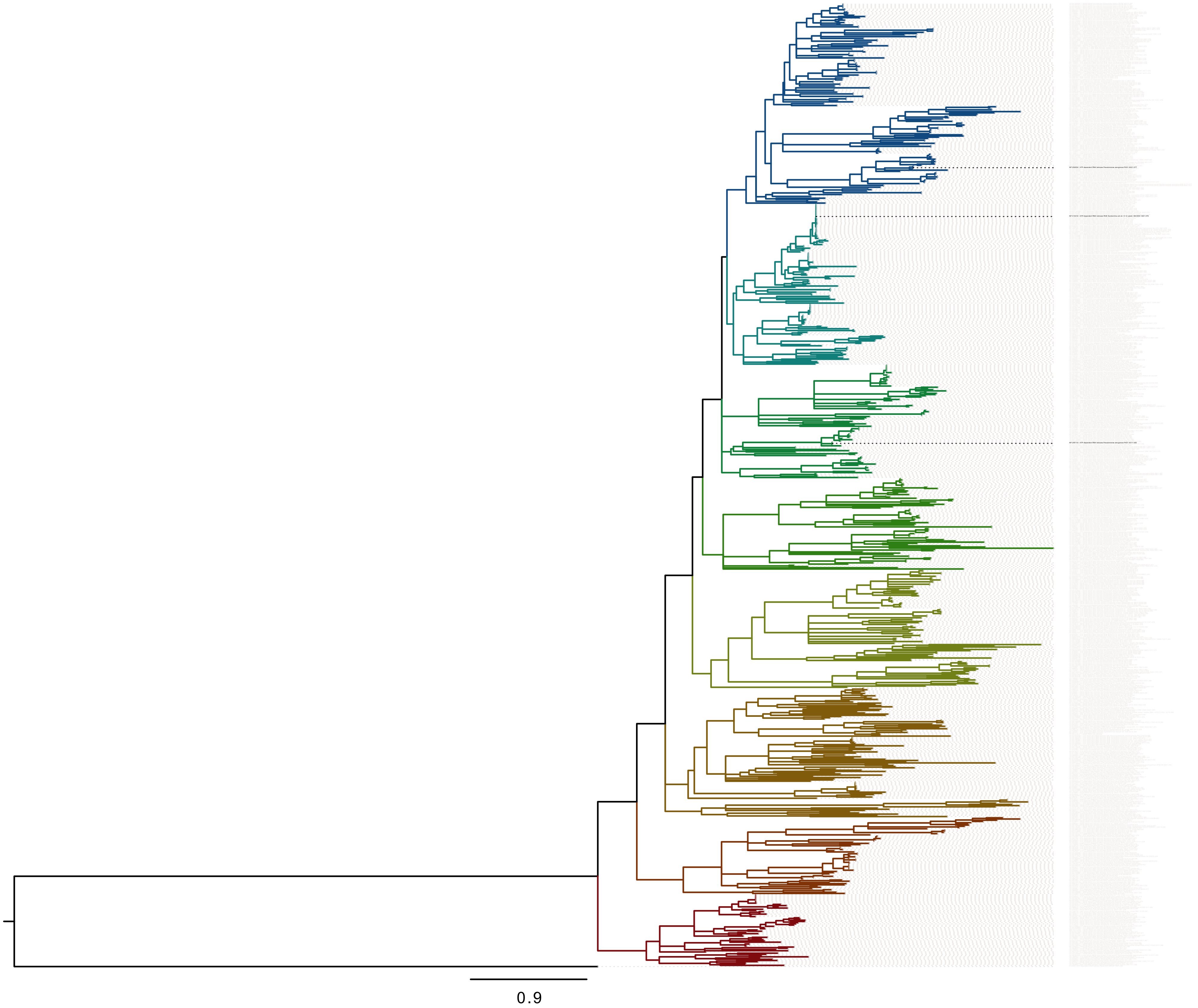
0.9

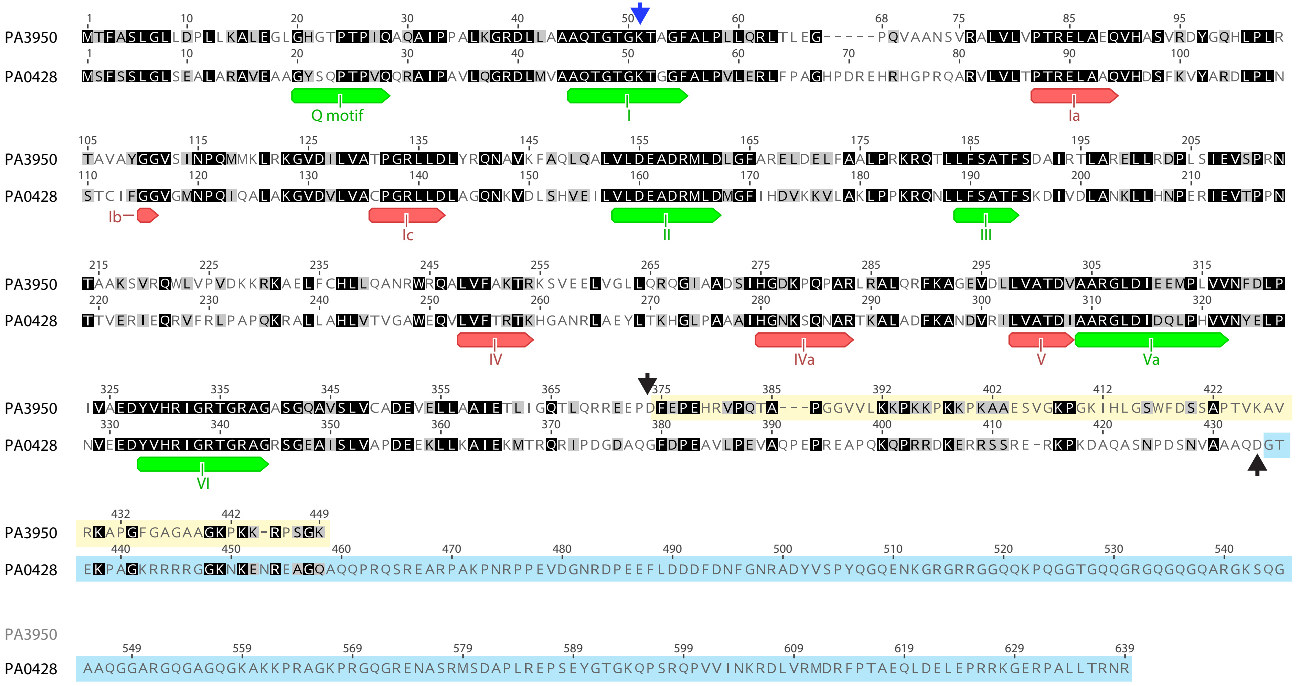

PA3950
PA0428
Q motif
I
Ia
Ib
Ic
II
III
IV
IVa
V
Va
VI

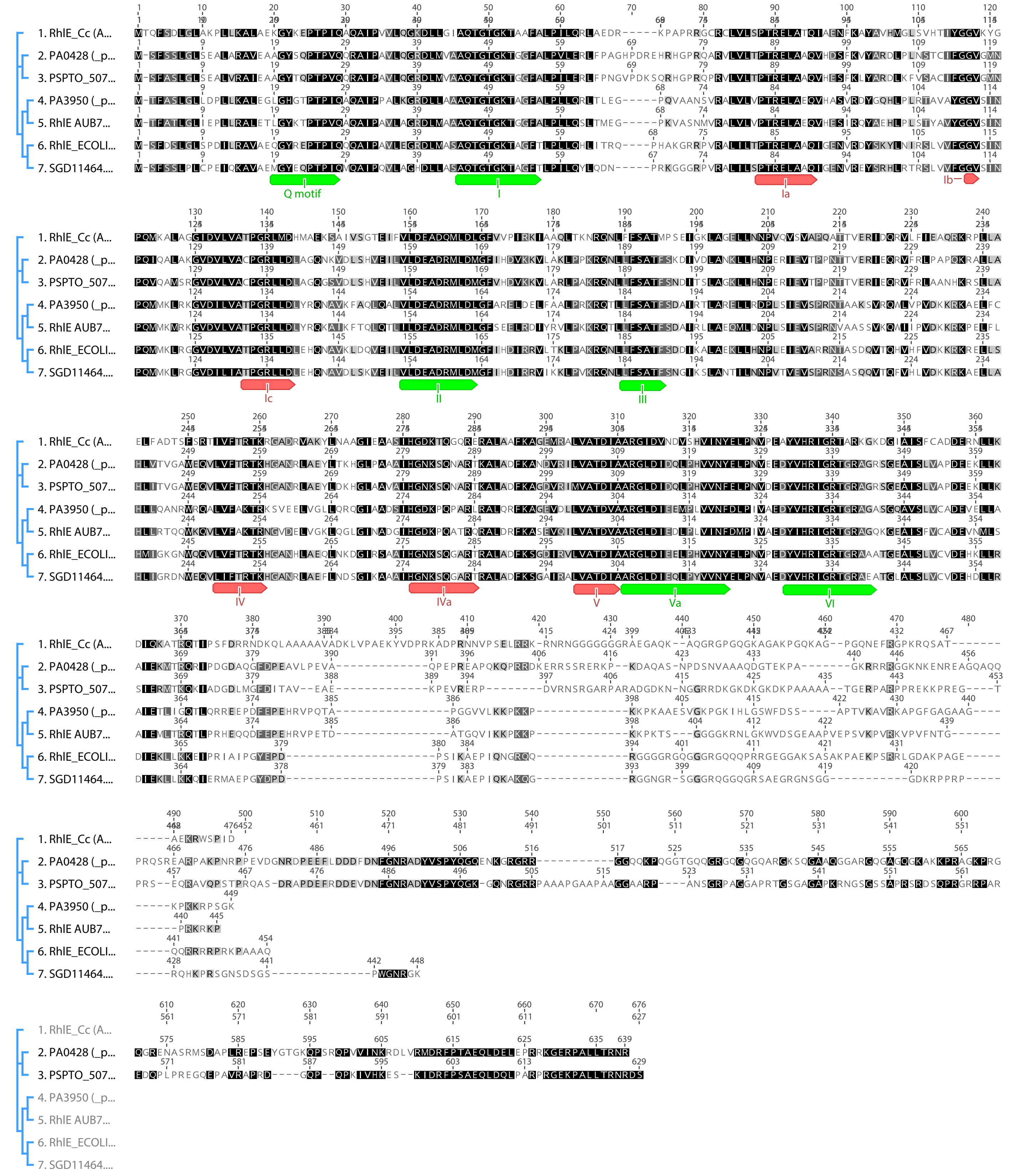

1. RhlE_Cc (A...
2. PA0428 (_p...
3. PSPTO_507...
4. PA3950 (_p...
5. RhlE AUB7...
6. RhlE_ECOLI...
7. SGD11464....
Q motif
I
Ia
Ib
Ic
II
III
IV
IVa
V
Va
VI

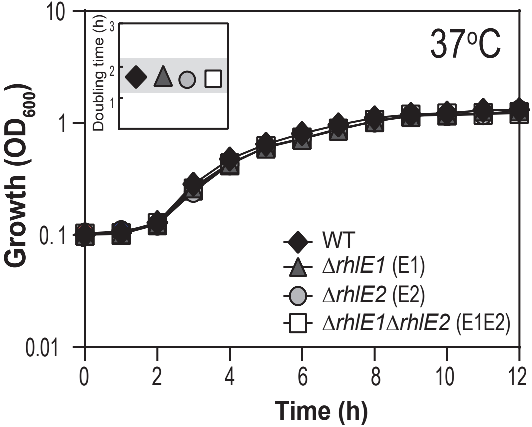
37oC
Doubling time (h)
3
2
1
Growth (OD600)
10
1
0.1
0.01
0
2
4
6
8
10
12
Time (h)
WT
ΔrhlE1 (E1)
ΔrhlE2 (E2)
ΔrhlE1ΔrhlE2 (E1E2)

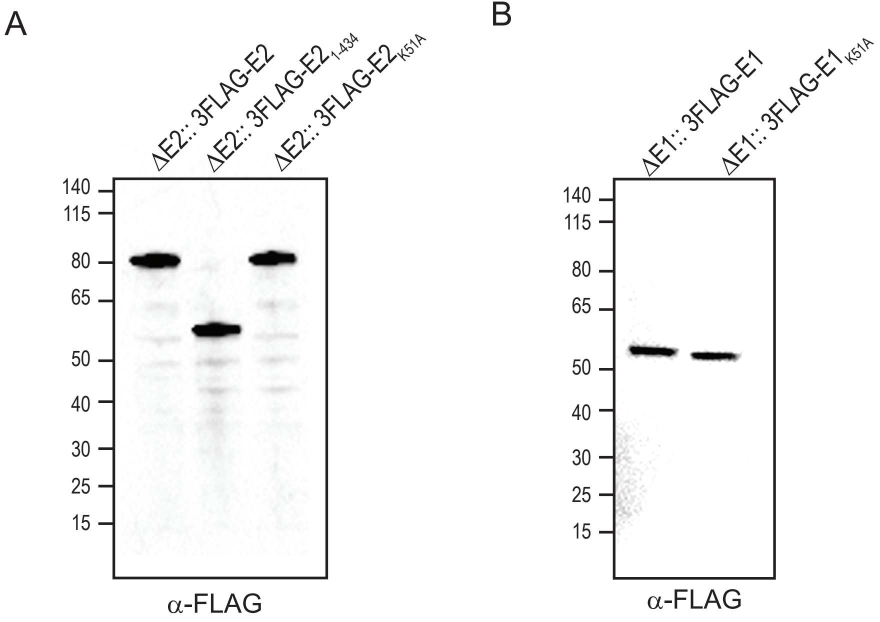
A
ΔE2:: 3FLAG-E2
ΔE2:: 3FLAG-E2$_{1\text{-}434}$
ΔE2:: 3FLAG-E2$_{K51A}$
140
115
80
65
50
40
30
25
15
α-FLAG
B
ΔE1:: 3FLAG-E1
ΔE1:: 3FLAG-E1$_{K51A}$
140
115
80
65
50
40
30
25
15
α-FLAG

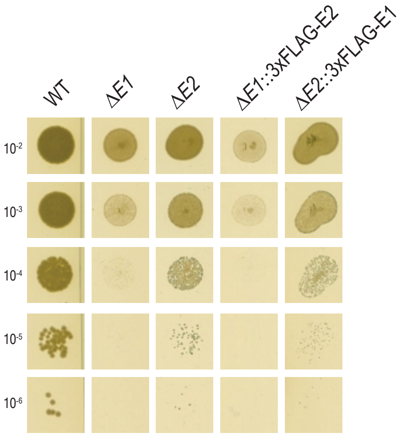
WT
ΔE1
ΔE2
ΔE1::3xFLAG-E2
ΔE2::3xFLAG-E1
10-2
10-3
10-4
10-5
10-6

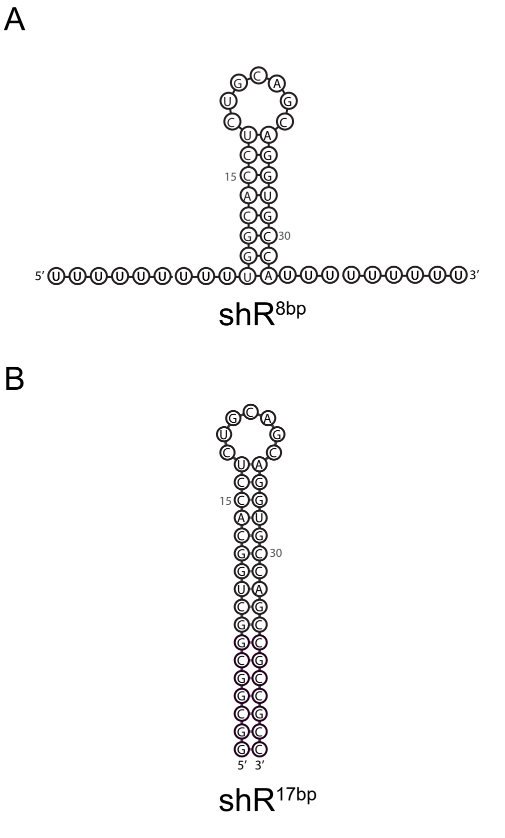
A
15
30
5′
3′
shR8bp
B
15
30
5′ 3′
shR17bp

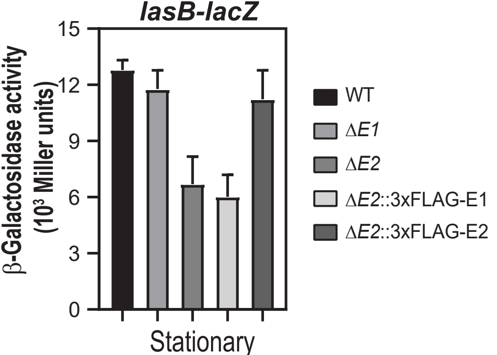
lasB-lacZ
β-Galactosidase activity
(10³ Miller units)
15
12
9
6
3
0
Stationary
WT
ΔE1
ΔE2
ΔE2::3xFLAG-E1
ΔE2::3xFLAG-E2

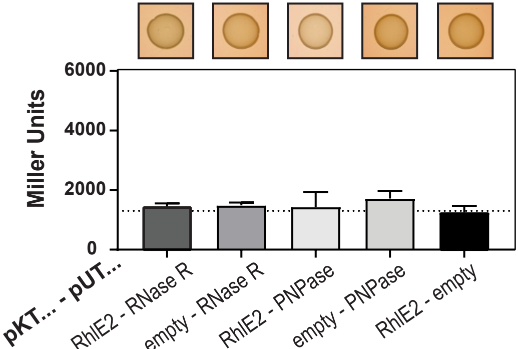

6000
4000
2000
0
Miller Units
pKT... - pUT...
RhlE2 - RNase R
empty - RNase R
RhlE2 - PNPase
empty - PNPase
RhlE2 - empty
